# Skipping without and with hurdles in bipedal macaque: Global mechanics

**DOI:** 10.1101/2023.08.26.554925

**Authors:** Reinhard Blickhan, Emanuel Andrada, Eishi Hirasaki, Naomichi Ogihara

## Abstract

Macaques trained to perform bipedally used running gaits across a wide range of speed. At higher speeds they preferred unilateral skipping (galloping). The same asymmetric stepping pattern was used while hurdling across two low obstacles placed at the distance of a stride within our experimental track. In bipedal macaques during skipping, we expected a differential use of the trailing and leading legs. The present study investigated global properties of the effective and virtual leg, the location of the virtual pivot point (VPP), and the energetics of the center of mass (CoM), with the aim of clarifying the differential leg operation during skipping in bipedal macaques. Macaques skipping displayed minor double support and aerial phases during one stride. Asymmetric leg use indicated by differences in leg kinematics. Axial damping and tangential leg work did not influence the indifferent peak ground reaction forces and impulses, but resulted in a lift of the CoM during contact of the leading leg. The aerial phase was largely due to the use of the double support. Hurdling amplified the differences. Here, higher ground reaction forces combined with increased double support provided the vertical impulse to overcome the hurdles. Following CoM dynamics during a stride skipping and hurdling represented bouncing gaits. The elevation of the VPP of bipedal macaques resembled that of human walking and running in the trailing and leading phases, respectively. Due to anatomical restrictions, macaque unilateral skipping differs from that of humans, and may represent an intermediate gait between grounded and aerial running.

## Introduction

In the wild, macaques prefer to locomote quadrupedally (Chatani, 2003; Fiers et al., 2013). The individuals trained at the Suo Monkey Performance Association, Japan, learned to post and locomote bipedally while guided on a leach. In this situation the macaques demonstrated their jumping ability traversing single hurdles of up to two meters height. They walked along the theater, but they never seemed to run with aerial phases. By investigating their running ability we discovered that macaques were able to use a variety of symmetrical gaits such as walking, grounded running (running gait without aerial phases), aerial running (with two aerial pases) and hopping (Ogihara et al., 2010; Ogihara et al., 2018). We also found that macaques preferred to bounce instead of vaulting over stiff legs at Froude speeds above 0.4 and used grounded running across a wide range of speed (Ogihara et al., 2018; Blickhan et al., 2018; Blickhan et al., 2021), even though humans usually avoid this gait bcause it is seemingly energetically more expensive than aerial running (Bonnaerens et al., 2018; Rummel et al., 2009). The compliant legs of macaques facilitated this gait (Andrada et al., 2020). Despite of some morphological adaptations to bipedal walking such as a human-like lordosis and more robust femora (Nakatsukasa et al., 2006), restricted hip joint extension (Ogihara et al., 2007) especially enforces crouched leg posture (Blickhan et al., 2021) and high leg compliance (Blickhan et al., 2018).

Like children of the age of about five (Roncesvalles et al., 2001), the bipedal macaques used unilateral skipping (galloping) when guided for fast locomotion (Ogihara et al., 2018). Skipping is a bouncing gait entailing a single aerial phase and a double support phase in one stride. Adult humans again usually avoid this gait but prefer the metabolically cheaper walk or run (Fiers et al., 2013). Nevertheless, like running, skipping can be self-stable and quite robust against disturbances (Andrada et al., 2016; Müller and Andrada, 2018).

In unilateral skipping or galloping, the same legs constitute the leading leg, i.e. the leg with ground contact which is followed by an aerial phase, and the trailing leg, i.e. the leg in contact after the aerial phase. In human skipping, the coordination at the hip joint seems to enforce a differential operation of the trailing and of the leading leg (Pequera et al. 2021). This was also expected in the macaque. We hypothesize, that the differential function of the legs observed in humans is also used by the macaques, despite of musculo-skeletal limitations.

The global properties of the legs ascertain skipping dynamics. This includes the angle of attack, leg stiffness and work done by the legs. Leg stiffness did not differ between grounded running and running (Blickhan et al., 2018). However, the contribution of a parallel damper shifted from damping to work. Also, the tangential contributions varied resulting in a shift from an elevated virtual pivot point (VPP: Maus et al., 2010) during grounded running to a human like pattern during aerial running. The global leg properties were adapted for each gait. This points towards the ability of differential leg operation in the macaque.

Compliant leg operation combined with limited leg extension could limit skipping ability. However, some bird species use skipping despite of compliant legs (Verstappen and Aerts, 2000; Alexander, 2004). Further, the macaques used with pleasure a skipping gait when they crossed hurdles placed at the distance of about a stride length. How does this influence the dynamics and kinematics of skipping? The present study aims to clarify the differential leg and trunk operation during skipping in bipedal macaques by analyzing global properties of the effective leg (between the center of pressure, CoP, and the hip) and virtual leg (between the CoP and the center of mass, CoM), the location of the VPP and the energetics of the CoM.

## Methods

More details of the methods largely repeated here for convenience are published in Ogihara et al., 2018) and Blickhan et al., 2018).

### Subjects

The macaques performed at the Suo Monkey Performance Association (Kumamoto, Japan). The three adult, male macaques (*Ku*, *Po*, *Fu*; age: 15, 13, 12 *years* mass: 8.64, 8.81, 8.79 *kg*) had been trained for bipedal walking and performances since the age of about one year. The grand means of leg lengths during the stance phase was 0.399, 0.339, 0.405 *m* for the effective leg, (*l*_0_) and 0.529, 0.465, 0.520 *m* for the virtual leg, (*l*_*c*0_) respectively. These were used for normalization.

### Setup

The macaques run across a flat wooden track (length: 5 m) with two embedded force plates (0.4 m x 0.6 m). During hurdling two hurdles (height: 0.1 m) were placed at the binning and the end of the two force plates (0.81 m apart; Fig. 1A). Kinematics and ground reaction forces were captured with an eight-camera infrared motion capture system (Oqus 3+, Qualisys, Göteborg, Sweden) and the force plates (EPF-S-1.5KNSA13; Kyowa Dengyo, Tokyo), respectively, at a rate of 200 Hz.

**Fig. 1.**
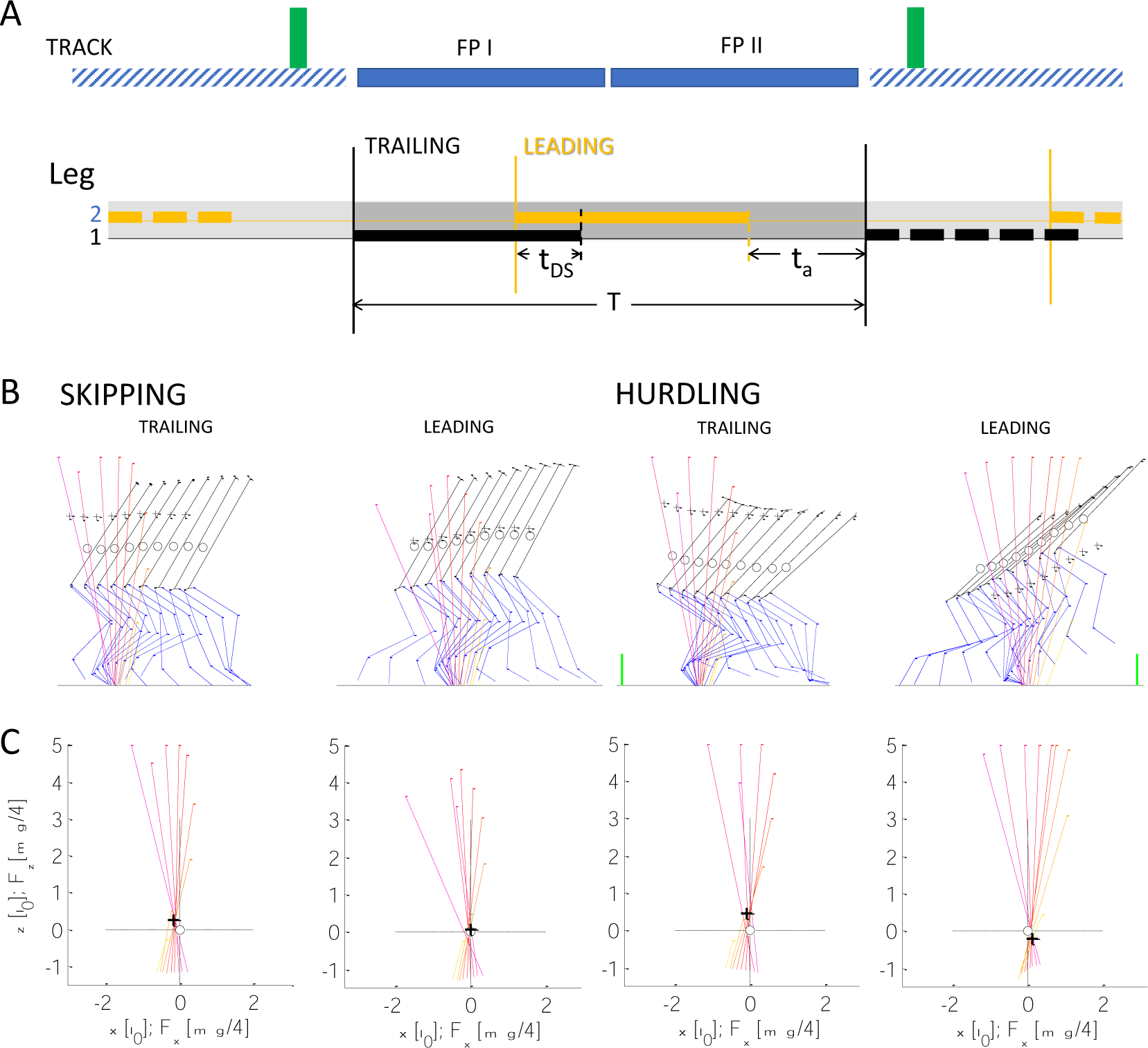
Kinematics and dynamics during skipping and hurdling. (A) Above: Setup. The macaques crossed a track with two force plates (FP I, FP II). During hurdling two hurdles (green squares) where placed before and after the force plates. Below: The stepping pattern during skipping and hurdling entails both a double support period (*t*_*DS*_ or *t*_*a*_ ≤ 0) and an aerial (*t*_*a*_ or *t*_*a*_ > 0) phase, defining the leading and trailing leg. The gait cycle usually starts with the touch down (solid vertical lines) of leg one, the trailing leg, onto the first force platform followed by the touch down of leg 2, the leading leg to the second force-plate with a double support until lift off (dashed vertical lines) of the trailing leg. The aerial period starts with the lift off of the leading leg. The stride ends with the touch down of the trailing leg after the flight phase. During 4 skipping trials the macaques stepped at the first platform with the leading leg. In these cases, the strides were defined by the consecutive touch downs of the leading leg. (B,C) Stick figures and VPP-Plot. In the stick figures two consecutive steps of a single trial are depicted, i.e. stick figures are overlapping (macaque: Fu). Circle: CoM, cross: VPP. (A) Stick-figures. Solid black line: trunk; blue solid line right leg; blue dashed line: left leg (note: different legs are used as trailing and leading legs in the two trials for skipping and hurdling); green lines within the two graphs during hurdling: hurdles (to scale). (B) VPP-graphs for trials depicted in (A). Contact times are divided in 8 segments. The color of the force vectors shifts with time from magenta to orange. Forces: 2 mg m^-1^. From left to right - speed: 2.2 m s^-1^, 2.2 m s^-1^, 1.7 m s^-1^, 1.7 m s^-1^: l_0_: 396 mm, 392 mm, 711 mm, 826 mm; floor lines: 647 mm, 889 mm, 711 mm, 826 mm; t_c_: 0.21 s, 0.22 s, 0.295 s, 0.255 s; ta: -0.015 s, 0.025 s, -0.035 s, 0.185 s; VPP_x_: -59 mm, -1 mm, -29 mm, 47 mm; VPP_z_: -107 mm, 24 mm, 173 mm, -87 mm.

### Procedure

An individual coach and caregiver guided the macaques across the track with a slack leash. A total of 15 markers were placed at the acromion, sternum xiphoid, tenth thoracic vertebra, anterior superior iliac spine, sacrum, greater trochanter, lateral epicondyle, lateral malleolus, and fifth metatarsal head.

### Ethical statement

The experiments were approved by the Animal Welfare and Animal Care Committee, Primate Research Institute, Kyoto University. All institutional guidelines were followed for this study. By rewards the macaques were easily motivated to walk bipedally. They were used to jump across high hurdles. Speed was freely selected and experiments were stopped as soon as signs of unwillingness surfaced.

### Data evaluation

Joint centers of the knee, the ankle, and the metatarsals were calculated by projecting the halve distance of the medial and lateral markers from the lateral markers perpendicular to the main plane of movement of the knee. The location of the hip was estimated as a projection perpendicular to the main plane of movement of the knee from the greater trochanter marker using a distance obtained from cadaver measurement (Ogihara et al., 2009). From this, the position of the segmental center of mass could be obtained using morphometric data (Ogihara et al., 2011). The center of mass of the trunk has been located on the line connecting mid hip joint (midpoint of the left and right joint centers) and mid shoulder. Within a presentation of the ground reaction forces with respect to the instantaneous CoM (CoM-fixed coordinate system) the VPP was calculated as the center of the waist (minimum horizontal width) established by the crossing of the extended ground reaction force vectors (first and last 10 % of contact time omitted; Blickhan et al., 2018).

In the present study, we focused on skipping and hurdling. Skipping was identified by a double support phase followed by an aerial phase (Fig. 1A). Both phases were decoded via the variable aerial phase, *t*_*a*_, with *t*_*a*_ ≤ 0 indicating double support and *t*_*a*_ > 0 indicating flight. The step and the leg in advance to the double support is termed “trailing”, and those in advance to the aerial phase was termed “leading”. In order to facilitate statistics as well as the analysis of the motion of the trunk segments, only sequences where a full data-set was available for both steps and no stumbling and distraction was observed were selected for further analysis. This selection resulted in a sample of 18 (Ku: 2; Fu: 8; Po: 8) strides for skipping an 31 (Ku: 4; Fu: 22; Po: 5) strides for hurdling. Global parameters where investigated during stance. The description of CoM data included a stride.

Froude speed was defined as 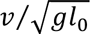 with g gravitational acceleration and *l*_0_, the mean of the touch down and take off length of the effective leg. The effective leg reaches from the CoP to the greater trochanter, and the virtual leg reaches from CoP to the CoM.

Leg stiffness, *k*, and damping, *D*, were calculated by fitting a parallel arrangement of a linear spring and a damper to the individual axial force-leg length data (*F*_*ax*_(*l*)) and we use a dimensionless formulation (Blickhan et al., 2018). These axial leg properties were complemented by tangential properties derived from leg torque-leg angle data (*M*(β_*leg*_) = *l* · *F*_*tan*_(β_*leg*_)). Axial, *W*_*ax*_, and tangential work, *W*_*tan*_, were calculated by the integrals with respect to leg length, *l*, and leg angle, β_*leg*_. The changes in potential energy, Δ*E*_*pot*_, were calculated from double integration of the ground reaction force.

The combined influence of leg (trailing vs. leading), Froude speed as a covariant, and of the individuals was tested general linear model (hierarchic -type I with repetitions; Bonferroni correction f = 141, IBM®SPSS®, Armonk, NY, U.S.A). The repetition refers to the steps of the leading and the trailing leg within the same stride (Tables 1, S1). This was complemented by univariate comparisons between skipping and hurdling considering the covariant Froude speed and the factor subject (Tables 1, S2).

Depending on normality of distribution (Lilliefors-test) parametric (t-test, unpaired t-test) or nonparametric tests (Wilkoxon sign-rank test, Wilcoxon) were performed (Table S3).

Custom software was written in MATLAB 14 (MathWorks, Natick, MA, U.S.A).

## RESULTS

### Global kinematics

The macaques used different leg angles in the trailing and leading period. Stride period, *T*, decreased with speed with individual variance (Fig. 3A; Table S1). Contact times, *t*_*c*_, were mostly shorter in the trailing leg than in the leading leg during hurdling (Fig. 3B; Table 1). The aerial time, *t*_*a*_, was used for classification. During skipping, an aerial phase (*t*_*a*_ > 0 s) follows a double support phase (*t*_*a*_ <= 0 s). The double support phase during skipping was always rather short (≥ −0.02 s; Fig. 3C). Trunk posture, β_*tru*_, showed a high interindividual variance (Figs. S1A, S2D,E; Table S1). In most cases β_*tru*_ decreased during stance. One subject (Fu) straightened in the trailing phase during hurdling. At lift off β_*tru*−*LO*_. was higher in the leading than in the trailing phase (Table 1), i.e. the subject was more erect. The leg lengthened during stance ((*l* − *l*_0_)_*TD*_ > (*l* − *l*_0_)_*LO*_; Figs. 2C, 3D,F; S1C). This was most pronounced in the leading leg while hurdling. There (*l* − *l*_0_)_*TD,LO*_ strongly differed in the trailing and leading phase (Table 1). Leg compression was most pronounced in the trailing leg while hurdling and the maximum compression was shifted towards midstance (Figs. 2C, S1C; Table 1). During hurdling leg rotation, β_*leg*_, was shifted towards a flatter angle of attack in the leading leg (Figs. 2B, 3H,I, S1B; β_*leg*−*TD,LO*_: Table 1).

**Fig. 2.**
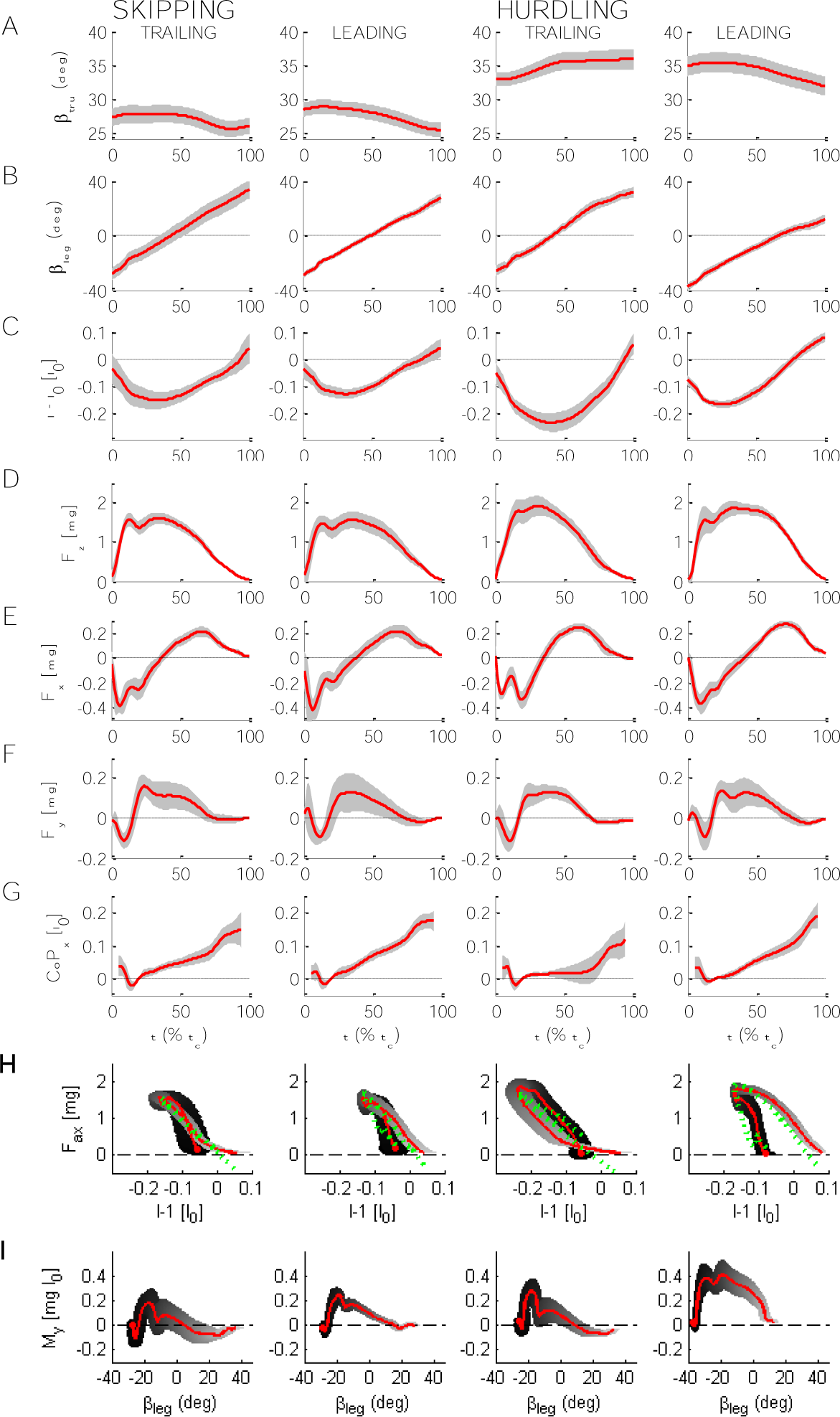
Global kinematic and dynamic parameters. Mean (red) and ±SD (shaded) of fime courses of global properties during skipping and hurdling in the trailing and the leading leg respectively. (A-G) *t*: time normalized to contact time, *t*_*c*_. (A) Trunk pitch, β_*tru*_. (B) Leg angle, β_*leg*_. (C) Change of leg length, (*l* − *l*_0_) *l*_0−1_. (D, E, F) Craniad, *F*_*z*_, anteriad, *F*_*x*_, and mediad, *F*_*y*_, components of ground reaction force. (G) Anteriad component of center of pressure, *CoP*_*x*_. *CoP*_*x*_ at 20% *t*_*c*_ set to 0 and the first and last 5% are omitted. (H) Axial loops, *F*_*ax*_(*l* − *l*_0_)*l*_0−1_. Green dashed lines: fittings based on Voigt-model. Filled circles: touch down. Shading from black at touch down to light gray at lift off. (I) Tangential loops, *M*_*y*_(β_*leg*_); filled circles: touch down; shading: from black at touch down to light gray at lift off. Mean contact times: 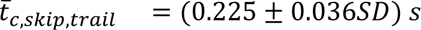; 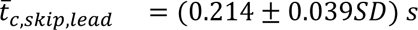; 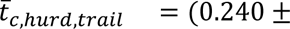; 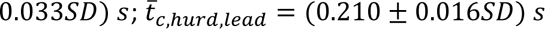. Mean leg length: 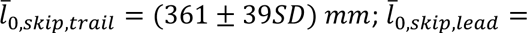; 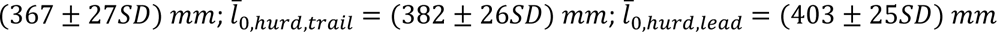.

**Fig. 3.**
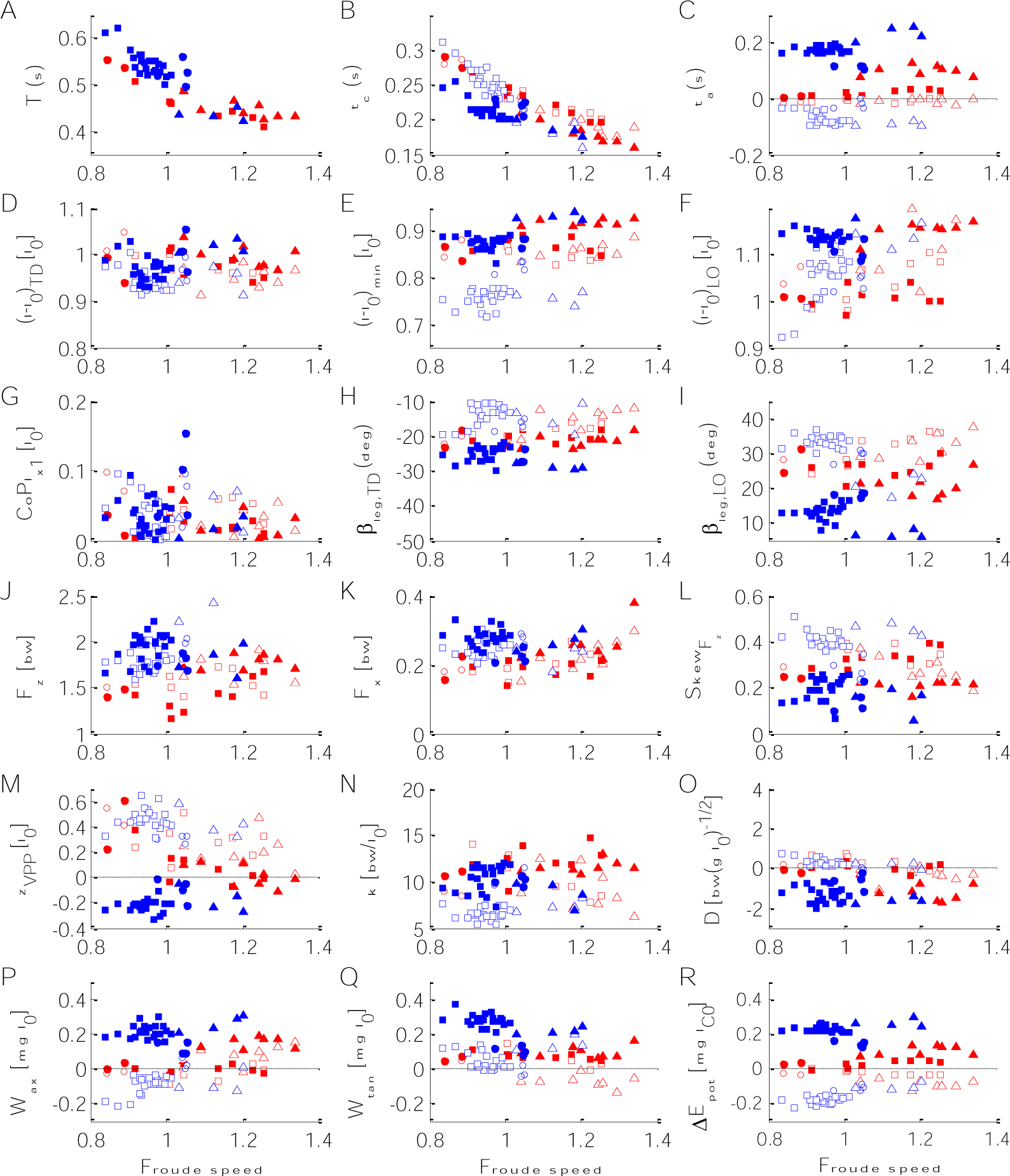
Dependence of kinematic, dynamic, and energetic stance parameters on Froude speed. Red: skipping; blue: hurdling; open: trailing leg; filled: leading leg. (A-C) Periods: (A) Stride period, *T*; (B) contact time, *t*_*c*_; aerial time, *t*_*a*_ (>0 flight; <=0 double support). (D-F) Lengthening of the effective leg from hip to CoP: (D) at touch down, (*l* − *l*_0_)_*TD*_; (E) at minimum length, (*l* − *l*_0_)_*min*_; (F) and lift off (*l* − *l*_0_)_*LO*_. (G) Length of CoP progression after 20% total length, *CoPl*_*x*1_. (H,I) Angle between leg and vertical: (H) at touch down, β_*leg,TD*_; (I) at lift off, β_*leg,LO*_. (J,K) Amplitude of the ground reaction force: (J) vertical force, *F*_*z*_; (K) anterior force, *F*_*x*_. (L) Skew of vertical force, *Skew*_*Fz*_. (M) Elevation of the VPP above the CoM, *z*_*VPP*_. (N) Stiffness of the leg, *k*. (O) Damping of the leg, *D*. (P-R) Work and energy: (P) axial work of the leg, *W*_*ax*_; (Q) tangential work, *W*_*tan*_; change in potential energy of the CoM, Δ*Pot*. Circle: Ku; square: Fu; triangle: Po. (Additional information: Fig.S2).

**Table 1.**
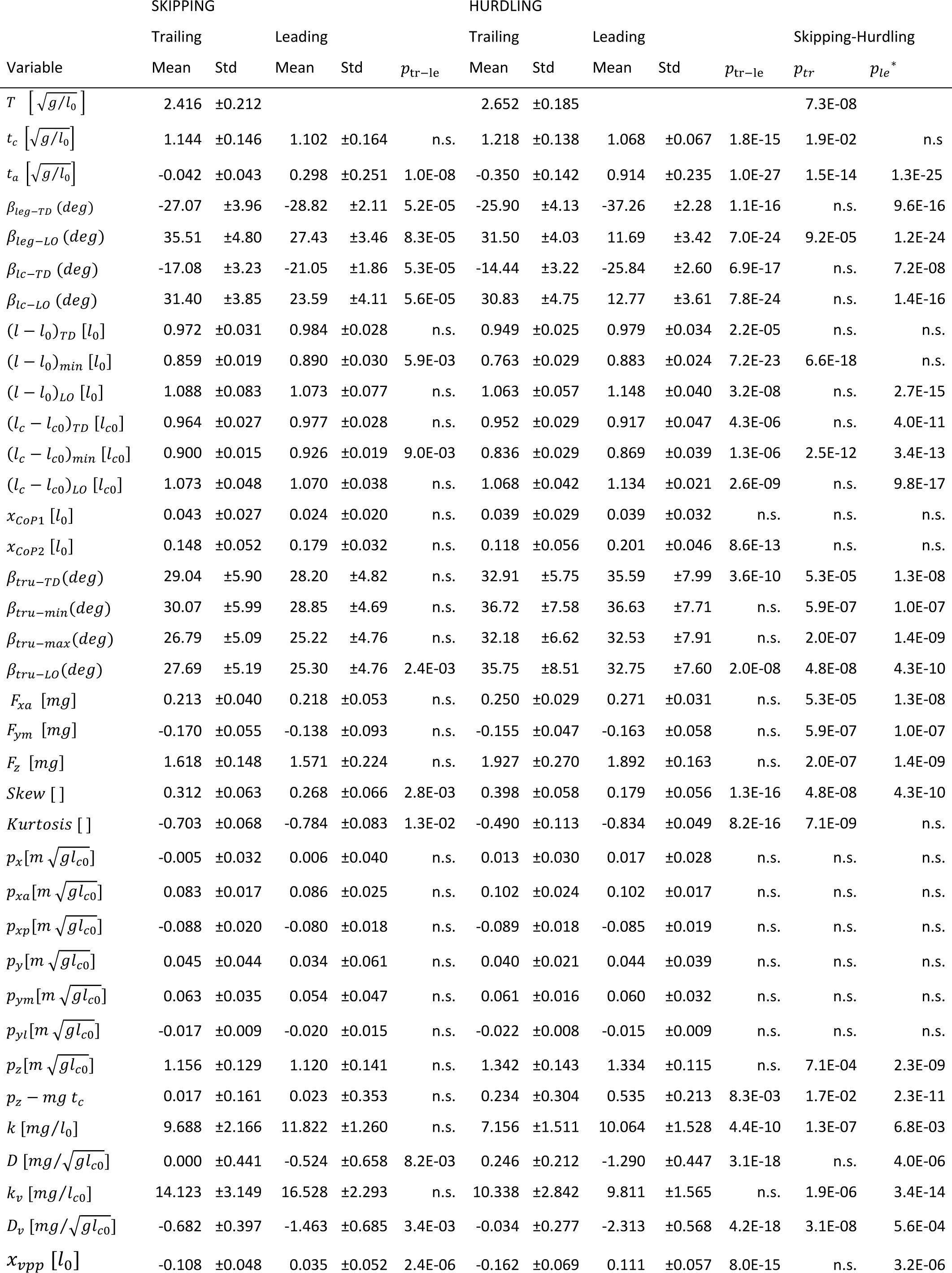
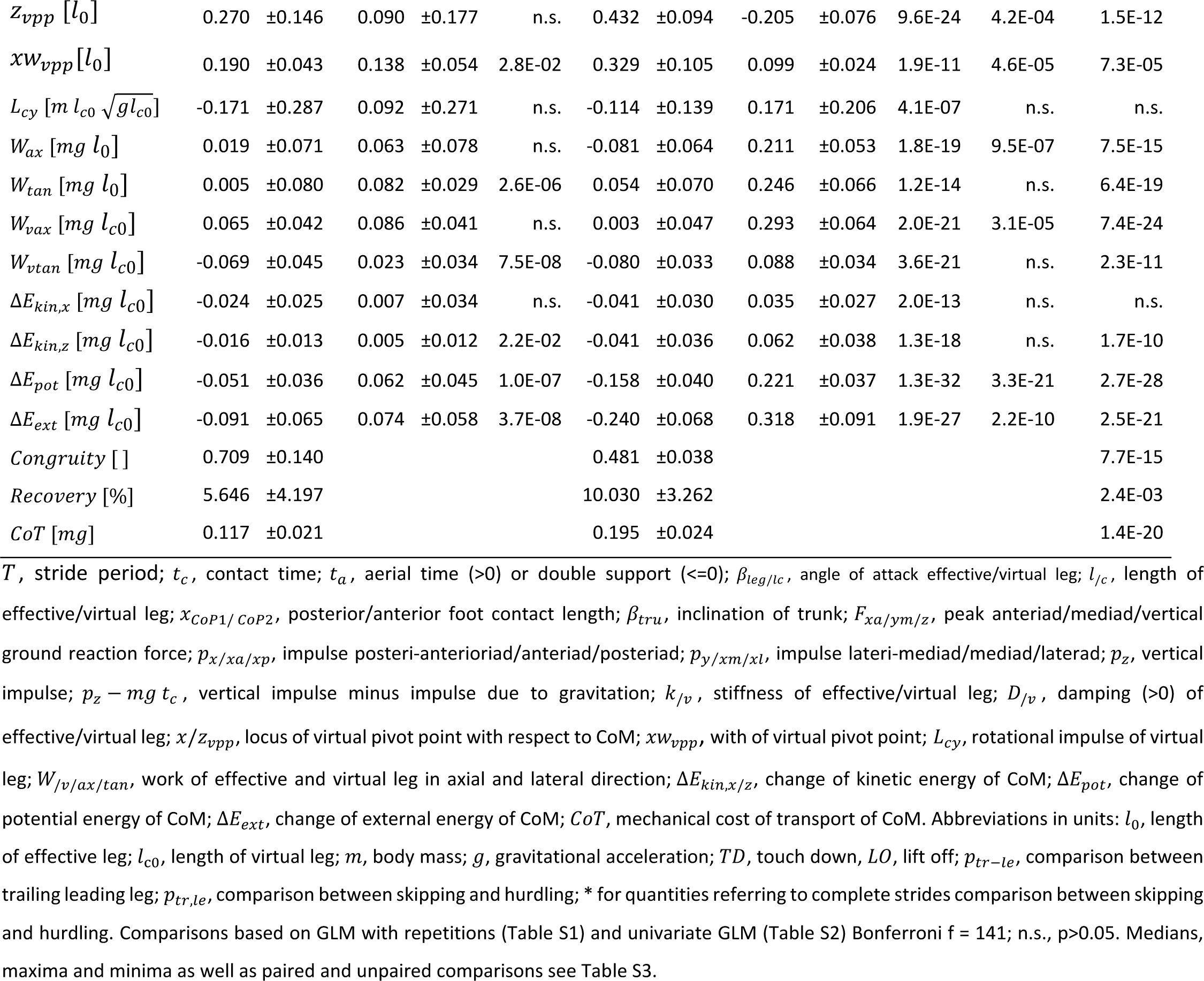
Comparison of global parameters between the leading and trailing leg and skipping and hurdling: timing, kinetics, leg properties, energetics.

### Forces, CoP and VPP

The time courses of the ground reaction forces, *F*_*x,y,z*_, were rather similar in the trailing and leading leg and during skipping and while crossing the hurdles (Fig. 2D,E,F). Nevertheless, skew of the vertical force, *F*_*z*_, was higher in the trailing than in the leading leg and the inverse was true for kurtosis (Figs. 2D, 3L, S1D; Table 1). The higher vertical force, *F*_*z*_, while jumping the hurdles were accompanied by increased propulsive forces, *F*_*xa*_ (Fig. 3J; Table 1). However, they did not differ in the leading and trailing leg (Table 1). The medial forces varied interindividually (Figs. S1F, S2F; Table S1). The progress of the CoP, *x*_*CoP*_, was reduced in the trailing leg during hurdling (Figs. 2G; 3G; S1G, Table 1, S2), which was most pronounced for a younger subject (Fu). The VPP was located posterior (*x*_*VPP*_ < 0) and above (*z*_*VPP*_ > 0) the CoM for trailing leg during both skipping and hurdling (Table 1). In the leading phase it was more focused (*xw*_*VPP*_ ; Table 1) and shifted towards the CoM (skipping) and anterior (*x*_*VPP*_ > 0) below (*z*_*VPP*_ > 0) the CoM during hurdling (Fig 1C; Fig. 3M). The horizontal shift of the VPP, *x*_*VPP*_, also the vertical shift, *z*_*VPP*_, during hurdling strongly differed in the trailing and leading period (Table 1).

### Leg stiffness, leg torque, and leg work

During hurdling stiffness, *k*, of the effective leg in the trailing phase was below the stiffness in the leading phase (sign. for hurdling; Fig. S1N; Table 1). The force-length-loops indicate energy absorption in the trailing leg during hurdling and positive axial work, *W*_*ax*_, in the leading leg (Fig. 2H; Table 1, S1). In the older subject (Ku) with its longer training history these differences were less pronounced (Figs. S1; S1H, S2P). The moment angle loops were positive in the leading leg both during skipping and hurdling being higher than the values observed in the trailing leg (*W*_*tan*_; Figs. 2I, 3Q, S1I; Table 1). The tangential Work, *W*_*tan*_, tended to decrease with Froude speed. The differences in potential energy of the CoM between lift off and touchdown, Δ*E*_*pot*_, indicated a lowering of the CoM in the trailing phase and a strong lift both during skipping and hurdling (Fig. S3; Table 1). Congruity was above 50% for skipping but about 50% for hurdling. Recovery was always below 20% during the stride (Table 1).

## Discussion

### Skipping

Walking, grounded and aerial running are lateral-symmetrical. In contrast skipping and hurdling are by definition lateral-asymmetrical. A differential operation of the leading and trailing legs was observed.

The aerial phase was not due to an elevated impulse generated at the leading leg but due to the double support phase. A difference between trailing and leading with respect to the peak ground-reaction force *F*_*x,y,z*_ was not observed (Table 1). The impulse, *p*_*x,y,z*_, generated by each leg did not differ even when considering for the vertical component the adverse action of gravity (−*mg* · *t*_*c*_). The more platykurtic form of the vertical force component, *F*_*z*_(*t*), of leading leg is compensated by its reduced contact time, *t*_*c*_. Positive skewness is reduced in the leading leg as compared to the trailing leg increasing the impulse during double support. The impulse generated by the combined action of both legs during the stride was sufficient to generate the short aerial phase after lift-off of the leading leg. The time course of the vertical velocity (Fig. S3) visualizes that the double support reduces the drop of the velocity. The double support was crucial to generate sufficient total impulse for the short aerial phase.

Differences with respect to timing were crucial for the gait. These differences were facilitated by differences in leg operation with respect to leg kinematics and leg compliance. The slightly lower minimum of the trailing leg length, (*l* − *l*_0_)_*min*_, indicates a higher leg compression (Fig. 2C, Table 1). A similar peak force at higher leg compression indicates a reduced leg stiffness, *k*, for the trailing leg. This is substantiated especially for hurdling by the fittings of the spring damper models (Fig. 2H; Table 1). In other words, the same peak force within a reduced contact time was generated in the leading leg with increased leg stiffness.

Differences in kinematics result in differential energetics. The trailing leg appears to operate quasi-elastic at the global level. In contrast, the leading leg lengthens and generates work (*W*_*ax*_; Table 1). The leg angle at touch down, β_*leg*−*TD*_, was flatter in the leading as compared to the trailing phase, it was steeper in the leading leg at lift off, β_*leg*−*LO*_ (Table 1). The flat angle of attack at lift off of the trailing leg facilitated the generation of the double support. The flat lift of angle, β_*leg*−*LO*_,of the trailing leg resulted in a lowering of the CoM in the trailing phase and in a reduction of potential energy, Δ*E*_*pot*_, of the CoM which was lifted in the leading phase (Table 1). The double support provided the impulse not only to stop the falling of the CoM in the trailing phase but to provide sufficient impulse to lift the CoM in the leading phase. Despite of higher vertical impulse, *p*_*z*_, as compared to horizontal, *p*_*x*_, the mixed terms result in higher fluctuations of the horizontal velocity. The fluctuations in kinetic energy were less than those of the potential energy (Fig. S3). The changes in external energy of the CoM, Δ*E*_*ext*_, were dominated by the contributions of the potential energy, Δ*E*_*pot*_. The dimensionless values of the latter correspond to the dimensionless changes in lift which are 5% *l*_*c*0_ or about 2.5 cm. The macaques had a smooth ride during skipping. Nevertheless, with ca. 70% congruity and ca. 6 % recovery skipping in macaques can be classified as a bouncing gait. The virtual leg did axial work, *W*_*vax*_ (Table 1) in agreement with the increase of total translational energy of the CoM. Assuring falling on their arms the net rotational impulses, *L*_*cy*_, generated by the ground reaction force with respect to the CoM is clockwise in the trailing phase and then reverse in the leading phase (n.s., Table 1). This is supported by the placement of the virtual pivot point behind, *x*_*VPP*_, and above, *z*_*VPP*_, the CoM in the trailing leg and very close to the CoM in the leading leg. In the latter, the ground reaction forces focused more precisely, *x*_*wpp*_, and closer to the CoM. The trunk was slightly more erect before lift of, β_*tru*−*LO*_, of the leading leg. The leading leg produces tangential work, *W*_*tan*,ℎ*ip*_, to compensate the rotational impulse (Table 1). The hip placement combined with the macaque’s posture enforces tangential work of the effective leg during retraction. Unfortunately, there remains an imbalance in our trials: The tangential work produced by the virtual leading leg, *W*_*vtan*_, is less than the absorption in the trailing phase (Table 1). This as well as the unbalanced rotational impulse, *L*_*cy*_, is also reflected in the placement of the virtual pivot point considering the two steps.

### Hurdling and differences to skipping

The differences between the kinetic parameters describing the trailing and leading steps were much more accentuated during hurdling (Tables 1, S2, S3).

Surprisingly, peak ground reaction force *F*_*x,y,z*_ and the impulses *p*_*x,y,z*_ during hurdling did not differ between the legs despite their enhanced value with respect to skipping (Table 1). However, the vertical impulse after considering gravity clearly differed between the legs for hurdling (Table 1). In the leading leg, the contact time, *t*_*c*_, was not shorter than during skipping, despite of slight lengthening of the contact during the trailing phase, and a lengthening in the stride period, *T* (Table 1). During skipping the double support was only marginal, but it was largely enhanced during hurdling. It was this enhanced double support providing the impulse to clear the hurdles. During hurdling as compared to skipping differences in form of the time courses of the vertical component of the ground reaction force of the trailing and leading leg were much more accentuated: During hurdling, left skewness was enhanced in the trailing leg and reduced in the leading leg, and kurtosis (excess) was reduced in the trailing leg and enhanced in the leading leg as compared to skipping (Table 1). Skewness indicates enhanced landing impacts after the flight phase. The leading leg was stiffer, *k*, than the trailing leg, i.e. as during skipping a lower compression (*l* − *l*_0_)_*min*_ within a reduced contact time resulted in similar forces (Table 1). As compared to skipping, the trailing leg was even more compliant (Table 1). A strong extension of the leading leg was observed at lift of (*l* − *l*_0_)_*LO*_ resulting in a high lift of the CoM, Δ*Epot*, during contact (Table 1). This was amplified by a rather steep angle of attack at lift off, β_*leg*−*LO*_ (Table 1). The leading leg was placed under a flat angle of attack, β_*leg*−*TD*_ and operated in a much more asymmetric mode as compared to the trailing leg. The leading leg produced axial work (*W*_*ax*,ℎ*ip*_; Table 1). This is expressed in the negative damping, D, parallel to the leg spring (Table 1). In the trailing leg the damper absorbed energy. The axial work of the virtual leg, *W*_*vax*_, was also concentrated on the leading leg (Table 1). The differences of the positions of the virtual pivot point, *x*, *z*_*VPP*_, were more accentuated as during skipping, indicating a more walking like trailing step and running like leading step (Table 1). As during skipping tangential work of the leg, *W*_*tan*_, was observed in both legs with clearly higher values in the leading phase. As in skipping, the imbalance in tangential work within the stride was reduced in the virtual leg (*W*_*vtan*_) as was also indicated by the differences of the generated rotational impulse (*L*_*cy*_; Table 1). The considerable tangential work, *W*_*tan*_, in the leading leg was counteracted by the erecting trunk (β_*tru*_; Table 1). The posterior placement of the hip enforces, in combination with the requirement to focus the forces to the CoM in preparation of the aerial phase, tangential work of the leg. The leg moment was counteracted by the movement of the trunk. Remarkably, the increased double support and the continued vertical acceleration of the CoM (Fig. S3) resulted in a congruity of about 50%, which is reduced as compared to skipping and is conceived as being the border between walking and running (e.g. Andrada et al., 2013).

By placing the hurdles before and after the two force plates, we provoked a movement pattern close to skipping. Without hurdles, skipping was the gait preferred by the macaques at higher Froude speeds (Ogihara et al., 2018). Skipping seemed to be convenient. It may have been this convenience why the macaues only rarely tried to run regularly. The fact that the macaques crossed the hurdles with ease proofs that they were able to generate higher flight phases using a similar bipedal rhythm. During skipping, there were differences with respect to the operation of the trailing and leading leg. However, these differences seem to represent only a minor deviation from standard mode of operation. As indicated above, the use of a double support phase, i.e. a rhythmical parameter, seemed to be essential to generate the short flight. The differential parameters were useful to cope with the consequences. During hurdling, these differences were largely exaggerated. Here, both kinematic and kinetic properties of the legs and its joints, as well as the role of the trunk strongly supported the different function of the trailing and leading leg. Two of the subjects had a preferred side (trailing leg for each individual: *left*/*right*, Ku: 6/0; Fu: 11/19; Po: 0/14).

### Comparison of skipping in human and macaque

Human parameters differ in some respect from those observed in macaques. and the magnifying observations during hurdling. The leg angles, β_*leg*_, with respect to the horizontal axis was less at the touch down of the trailing and at the lift off of the leading leg during skipping (macaque skip / hurdle / human skip, β_*leg,trail*−*TD,TO*_: 62.9, 125.5 / 64.1, 121.5 / 82.1, 122.2 (*deg*) ; β_*leg,lead*−*TD,LO*_: 61.2, 117.4 / 52.7, 101.7 / 57.4, 103.1 (*deg*) ; Müller and Andrada, 2018). Leg lengthening, Δ*l* = *l*_*LO*_ − *l*_*TD*_, is more expressed in the macaque (Δ*l* _*trail*_: 0.11 / 0.11 / −0.01 [*l*_0_]; Δ*l* _*lead*_: 0.09 / 0.17 / 0.025 [*l*_0_]; Müller and Andrada, 2018). Lift of the CoM, Δ*z*_*CoM*_[*l*_0_] = Δ*E*_*pot*_[*mgl*_0_], is less for skipping in macaques but higher for hurdling as compared to human skippers (Δ*z*_*CoM,trail*_: -0.05 / -0.16 / 0.1 [*l*_0_]; Δ*z*_*CoM,trail*_: 0.06 / 0.22 / 0.09 [*l*_0_]; Müller and Andrada, 2018), but in all cases the CoM is lowered in the trailing an lifted in the leading phase. The macaque used in general similar leg movements but with lower leg stiffness (see below).

As in macaques, unilateral skipping amplitudes of the vertical component of the ground-reaction force (*F*_*z,trail*_ = 2.28 [mg], *F*_*z,lead*_ = 2.14 [mg]) and its impulse (*p*_*z,trail,lead*_ = 2.09 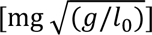) did not differ significantly between the leading and the trailing leg in human (Fiers et al., 2013). However, the amplitudes by far exceeded even the values observed in macaques during hurdling and so do the impulses in forward direction (*p*_*x,trail*_ = −0.030 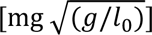; *p*_*x,lead*_ = 0.033 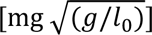. In both species, the trailing leg is decelerating and the leading leg is accelerating. However, the decelerations and accelerations as related to the vertical impulse were in the macaque less than 20% of the contributions during human skipping. During hurdling in macaques, both legs were accelerating and the ratios between horizontal and vertical impulses reached almost the human values. The lower oscillations of the horizontal energy in the macaque during skipping seems to be a matter of convenience. There was net acceleration in our trials during hurdling, the track allowed about three strides. The human subjects preferred to locomote on a treadmill at about the same Froude speed (*Fr* = 1.06 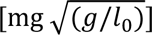 but with a considerable shorter stride duration (*T* = 2.06 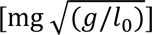. Correspondingly, the contact times were shorter (*t*_*c,trail*_ = 0.76 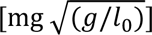; *t*_*c,lead*_ = 0.78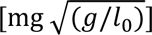). Human bipedal gallopers used a higher leg stiffness as compared to macaques. This is also confirmed in a recent study where kinematic parameters were used to estimate leg stiffness during unilateral skipping or galloping (Pequera et al., 2021). In this study, the stiffness of the trailing leg by far exceeded the values obtained for the leading leg and the values obtained in our study for the macaques (from regression Fr = 1: *k*_*trail*_ = 46.4 [*mg*⁄*l*_0_]; *k*_*lead*_ = 23.8 [*mg*⁄*l*_0_], Pequera et al., 2021). In our study in human unilateral skipping, stiffness of the leading leg was enhanced as in the macaque (*Fr* ≈ 1; *k*_*trail*_ = 34.9 [*mg*/*l*_0_]; *k*_*lead*_ = 44.1 [*mg*⁄*l*_0_]; Müller and Andrada, 2018). The macaques skip with more compliant legs.

For the trailing leg the direction of the ground reaction force as quantified by *z*_*VPP*_ for skipping resembled the values found for the macaques during running (0.20 [*l*_0_]), and for hurdling the values found during grounded running (0.38 [*l*_0_]; Blickhan et al., 2018). Both are within the range found during human walking (Maus et al., 2010; Vielemeyer et al., 2019). In the leading leg, the values move closer to the CoM and for hurdling even below the CoM. This resembles human running (Maus et al., 2010). At high speeds during human running the values are below the CoM (Drama and Badri-Spröwitz, 2020). Simulations demonstrate (Drama and Badri-Spröwitz, 2019) that the VPP in the vicinity of the CoM as found during slow human running facilitates exchange of energy between trunk and legs. The horizontal displacement of the VPP, *x*_*VPP*_, moves with increased trunk flexion posterior during human walking (Müller et al., 2017). In the macaques the different location of the VPP in the leading and trailing leg correlates with the transmitted rotational impulse *L*_*cy*_. However, the differences between trailing and leading leg are not significant during skipping. Nevertheless, whole body rotational impulse seems to be modified to provide secure landing in the trailing period and may also support the take off in the leading phase. During hurdling, the influence of the extended double support may affect the location of the VPP. During human walking, *z*_*VPP*_ drops to zero during the double support and looses focusation (Vielemeyer et al., 2021). This may help to adjust rotational moments and posture from step to step.

The energy of the CoM from touch down to take off in humans indicates clear horizontal acceleration in the trailing leg and deceleration the leading leg (Δ*E*_*kin,x,trail*_ = 0.098 [*mgl*_*c*0_]; Δ*E*_*kin,x,lead*_ = −0.084 [*mgl*_*c*0_]; Fiers et al., 2013), a vertical deceleration in the trailing and an acceleration in the leading leg (Δ*E*_*kin,z,trail*_ = −0.044 [*mgl*_*c*0_]; Δ*E*_*kin,z,lead*_ = 0.037 [*mgl*_*c*0_]), and a lowering of the CoM in the trailing and a lift in the leading leg (Δ*E*_*pot,trail*_ = −0.110 [*mgl*_*c*0_]; Δ*E*_*pot,lead*_ = 0.141 [*mgl*_*c*0_]). Such a pattern has also been documented in a trial in the pioneering study on bilateral skipping of Minetti et al. (Minetti, 1998; *Fr* = 0.84 ; Δ*E*_*kin,x,trail*_ = 0.078 [*mgl*_*c*0_]; Δ*E*_*kin,x,lead*_ = −0.046 [*mgl*_*c*0_] ; Δ*E*_*kin,z,trail*_ = −0.040 [*mgl*_*c*0_]; Δ*E*_*kin,z,lead*_ = 0.032 [*mgl*_*c*0_] ; Δ*E*_*pot,trail*_ = −0.121 [*mgl*_*c*0_]; Δ*E*_*pot,lead*_ = 0.128 [*mgl*_*c*0_]), and in the early study of Caldwell and Whitall (Caldwell and Whitall, 1995; *Fr* ≈ 1; Δ*E*_*kin,trail*_ ≈ 0.05 [*mgl*_*c*0_]; Δ*E*_*kin,lead*_ ≈ −0.06 [*mgl*_*c*0_]; Δ*E*_*pot,trail*_ ≈ −0.07 [*mgl*_*c*0_]; Δ*E*_*pot,lead*_ ≈ 0.1 [*mgl*_*c*0_]). This deviates from the pattern found in the macaque: in the macaque the horizontal energy decreased in the trailing leg and increased in the leading leg and the fluctuations of the kinetic energies were reduced during skipping. Human unilateral skippers also lower the CoM in the trailing phase to used a flatter angle of attack of the leading leg to redirect the horizontal kinetic energy gained in the trailing phase to generate lift for the flight. As in the macaques, according to the energetics of the CoM, skipping steps in humans were of the running type. There was no exchange between potential and kinetic energy or an inverted pendulum (comp. discussion in Fiers et al., 2013). In contrast to Minetti (1998), Pavei et al., 2015) documented recovery values for bilateral skipping close to running values (Fr = 1.01; recovery = 21 %). They are even lower for the skipping and hurdling macaques. The congruity values for skipping macaques were rather similar to the values obtained in the macaques during grounded and aerial running (Ogihara et al., 2018). Unilateral skipping represents an intermediate gait between grounded and aerial running. (The extended double support during hurdling modified the energetics of CoM.) The external mechanical cost of transport, *CoT*, observed in the macaques during skipping were of similar magnitude as the values observed during fast bilateral skipping in humans (0.08 < *CoT* [*mg*] < 0.25; Minetti, 1998). The values observed during hurdling were still less than the highest of humans during walking. For unilateral skipping (*CoT* ≈ 0.1 [*mg*], Caldwell and Whitall, 1995; 0.17 [*mg*], Fiers et al., 2013) similar and higher values are documented. Macaques avoided a bumpy ride.

In human locomotion, skipping is more expensive than running. If we assume this for the macaques then why did they prefer this gait? One reason could be stability. In the numerical simulation the point of operation (β_*vleg,trail*−*TD*_ = 72.9 *deg* ; β_*leg,lead*−*TD*_ = 68.9 ; *k*_*v,trail*_ = 14.1 [*mg*⁄*l*_*c*0_] ; *k*_*v,lead*_ = 16.5 [*mg*⁄*l*_*c*0_]) is close to but outside the selfstable region for a skipper with purely elastic legs (Andrada et al., 2016). However, this ignores the possibly stabilizing influence of the force generation (negative damping) parallel to the spring (*D*_*v,trail*_ = −0.68 [*mg*⁄(*gl*_*c*0_)^1/2^]; *D*_*v,lead*_ = −1.46 [*mg*⁄(*gl*_*c*0_)^1/2^]). The mechanical cost of transport of the CoM was less for grounded running and slightly less for running but much higher for hurdling (*CoT*_*GR*_ = 0.074 ± 0.010*SD* [*mg*], *p*_*GR,sk*_ = 5*E* − 9, *p*_*GR*,ℎ*u*_ = 5*E* − 12, *n*_*GR*_ = 38 ; *CoT*_*R*_ = 0.101 ± 0.013*SD* [*mg*] ; *p*_*R*,*sk*_ = 0.0020, *p*_*GR*,ℎ*u*_ = 1*E* − 13, *n*_*GR*_ = 46). We did not measure oxygen consumption. However, as in human locomotion with respect to mechanics skipping seems to be less convenient than symmetrical gaits. Despite of bipedal training, our macaques prefer quadrupedal locomotion possibly due to its reduced energetic cost (Nakatsukasa et al., 2004, Nakatsukasa et al., 2006). During fast quadrupedal locomotion they prefer a transverse gallop with a dominant hindlimb contribution (Kimura, 1992). Unilateral skipping or bipedal galloping represents a transverse gallop without forelimbs. Skipping may represent a preferred motor pattern for the bipedal macaques. Recent findings indicate that quadrupeds walk and trot with VPPs above the hip and the scapula (Andrada et al., 2023). It remains thus intriguing, if the differences in VPP heights depicted here between trailing and leading limbs holds for quadrupedal gallop, or they represent an adaptation to bipedal skipping.

## Acknowledgements

We express our gratitude to all the staff of the Suo Monkey Performance Association for their generous collaborations in these experiments. Thanks are due to Naoki Kitagawa, Kohta Ito, Hideki Oku, Mizuki Tani from Keio University, Yokohama, Japan, and Martin Götze Friedrich-Schiller University, Jena, Germany, for helping us during data collection. We also thank Ryoji Hayakawa, ArchiveTips, Inc., Tokyo, Japan, for support and preprocessing with Qualisys. The authors would like to thank Keio University for the guest professorship appointment to R.B. strongly facilitating this line of research.

## Author contributions

Conceptualization: R.B., N.O.; Methodology: E.A., N.O.; Software: R.B.; Formal analysis: R.B.; Investigation: R.B., E.H., N.O.; Resources: E.H., N.O.; Data curation: N.O.; Writing - original draft: R.B.; Writing - review & editing: E.A., N.O.; Supervision: N.O.; Project administration: N.O.; Funding acquisition: N.O, R.B., E.A..

## Data availability

Dynamic and kinematic data are available from figshare: DOI 10.6084/m9.figshare.24008052. Software for data processing are available on request from the corresponding author.

## Supplement

### TABLES SUPPLEMENT

**Table S1.** Comparison of global parameters between the leading and trailing leg for skipping and hurdling: timing, kinetics, leg properties, energetics. (Probabilities of GLM with repetitions.)

| Variables | SKIPPING |  |  |  |  | HURDLING |  |  |  |  |
| --- | --- | --- | --- | --- | --- | --- | --- | --- | --- | --- |
|  | tr-le | tr-le*Fr | tr-le*ani | Fr | ani | tr-le | tr-le*Fr | tr-le*ani | Fr | ani |
| $t_c [\sqrt{g/l_0}]$ | n.s. | n.s. | n.s. | 9.5E-10 | n.s. | 1.8E-15 | 1.5E-08 | 4.3E-02 | 2.4E-08 | n.s. |
| $t_a [\sqrt{g/l_0}]$ | 1.0E-08 | 9.9E-04 | n.s. | 7.1E-04 | 9.9E-04 | 1.0E-27 | 1.5E-05 | 1.6E-09 | 7.1E-02 | n.s. |
| $\beta_{leg-TD} (deg)$ | 5.2E-05 | 3.5E-02 | n.s. | n.s. | n.s. | 1.1E-16 | n.s. | n.s. | n.s. | n.s. |
| $\beta_{leg-LO} (deg)$ | 8.3E-05 | n.s. | n.s. | n.s. | n.s. | 7.0E-24 | 1.1E-03 | 3.2E-03 | 9.0E-08 | 9.4E-08 |
| $\beta_{lc-TD} (deg)$ | 5.3E-05 | 3.1E-02 | n.s. | n.s. | n.s. | 6.9E-17 | n.s. | n.s. | n.s. | n.s. |
| $\beta_{lc-LO} (deg)$ | 5.6E-05 | n.s. | n.s. | n.s. | n.s. | 7.8E-24 | 2.5E-03 | 1.8E-03 | 4.4E-08 | 2.7E-07 |
| $(l-l_0)_{TD} [l_0]$ | n.s. | n.s. | n.s. | n.s. | n.s. | 2.2E-05 | n.s. | n.s. | n.s. | n.s. |
| $(l-l_0)_{min} [l_0]$ | 5.9E-03 | n.s. | n.s. | n.s. | n.s. | 7.2E-23 | n.s. | 1.8E-07 | n.s. | n.s. |
| $(l-l_0)_{LO} [l_0]$ | n.s. | n.s. | n.s. | 5.6E-04 | 9.7E-03 | 3.2E-08 | n.s. | n.s. | 3.1E-08 | 2.3E-03 |
| $(l_c-l_{c0})_{TD} [l_{c0}]$ | n.s. | n.s. | n.s. | 2.8E-02 | 6.8E-03 | 4.3E-06 | 1.4E-04 | n.s. | 4.3E-02 | 3.7E-03 |
| $(l_c-l_{c0})_{min} [l_{c0}]$ | 9.0E-03 | n.s. | n.s. | n.s. | 2.7E-02 | 1.3E-06 | 2.8E-03 | n.s. | 6.2E-05 | 5.6E-04 |
| $(l_c-l_{c0})_{LO} [l_{c0}]$ | n.s. | n.s. | n.s. | 1.4E-02 | n.s. | 2.6E-09 | n.s. | n.s. | 1.4E-04 | 2.2E-02 |
| $x_{CoP1} [l_0]$ | n.s. | n.s. | n.s. | n.s. | n.s. | n.s. | n.s. | n.s. | n.s. | n.s. |
| $x_{CoP2} [l_0]$ | n.s. | n.s. | n.s. | n.s. | n.s. | 8.6E-13 | n.s. | n.s. | 7.1E-04 | 5.6E-05 |
| $\beta_{tru-TD} (deg)$ | n.s. | n.s. | n.s. | n.s. | 5.5E-03 | 3.6E-10 | 1.3E-05 | 1.2E-06 | 1.2E-09 | 2.6E-08 |
| $\beta_{tru-min} (deg)$ | n.s. | n.s. | n.s. | n.s. | n.s. | n.s. | n.s. | n.s. | 3.2E-08 | 1.3E-05 |
| $\beta_{tru-max} (deg)$ | n.s. | n.s. | n.s. | n.s. | n.s. | n.s. | 2.4E-07 | n.s. | 5.6E-12 | 1.3E-09 |
| $\beta_{tru-LO} (deg)$ | 2.4E-03 | n.s. | n.s. | n.s. | n.s. | 2.0E-08 | n.s. | n.s. | 2.2E-11 | 3.0E-07 |
| $F_{xa} [mg]$ | n.s. | n.s. | n.s. | n.s. | n.s. | n.s. | n.s. | n.s. | n.s. | n.s. |
| $F_{ym} [mg]$ | n.s. | n.s. | n.s. | n.s. | 5.7E-07 | n.s. | n.s. | n.s. | 2.1E-06 | 6.8E-06 |
| $F_z [mg]$ | n.s. | n.s. | n.s. | 2.6E-02 | n.s. | n.s. | 2.8E-04 | n.s. | 1.2E-09 | 4.4E-02 |
| *Skew* [ ] | 2.8E-03 | n.s. | 2.1E-03 | n.s. | 2.1E-03 | 1.3E-16 | n.s. | 6.1E-03 | n.s. | n.s. |
| *Kurtosis* [ ] | 1.3E-02 | n.s. | n.s. | n.s. | n.s. | 8.2E-16 | n.s. | n.s. | n.s. | n.s. |
| $p_x[m\sqrt{gl_{c0}}]$ | n.s. | n.s. | n.s. | n.s. | n.s. | n.s. | n.s. | n.s. | n.s. | n.s. |
| $p_{xa}[m\sqrt{gl_{c0}}]$ | n.s. | n.s. | n.s. | n.s. | n.s. | n.s. | n.s. | n.s. | n.s. | n.s. |
| $p_{xp}[m\sqrt{gl_{c0}}]$ | n.s. | n.s. | n.s. | n.s. | n.s. | n.s. | n.s. | n.s. | n.s. | n.s. |
| $p_y[m\sqrt{gl_{c0}}]$ | n.s. | n.s. | n.s. | 1.4E-03 | 3.3E-06 | n.s. | n.s. | n.s. | 5.1E-03 | 2.1E-05 |
| $p_{ym}[m\sqrt{gl_{c0}}]$ | n.s. | n.s. | n.s. | n.s. | n.s. | n.s. | n.s. | n.s. | n.s. | 5.1E-06 |
| $p_{yl}[m\sqrt{gl_{c0}}]$ | n.s. | n.s. | n.s. | 1.1E-02 | 4.2E-04 | n.s. | n.s. | n.s. | 3.8E-03 | 7.8E-03 |
| $p_z[m\sqrt{gl_{c0}}]$ | n.s. | n.s. | n.s. | 7.1E-04 | 6.0E-06 | n.s. | n.s. | n.s. | n.s. | n.s. |
| $p_z - mg t_c [m\sqrt{gl_{c0}}]$ | n.s. | n.s. | n.s. | 2.3E-05 | 1.4E-04 | 8.3E-03 | 3.0E-03 | n.s. | 1.4E-04 | n.s. |
| $k [mg/l_0]$ | n.s. | n.s. | n.s. | n.s. | n.s. | 4.4E-10 | 1.8E-02 | 1.1E-02 | n.s. | 3.3E-02 |
| $D [mg/\sqrt{gl_{c0}}]$ | 8.2E-03 | n.s. | n.s. | n.s. | n.s. | 3.1E-18 | n.s. | 1.1E-03 | n.s. | n.s. |
| $k_v [mg/l_{c0}]$ | n.s. | n.s. | n.s. | n.s. | n.s. | n.s. | n.s. | 1.2E-02 | n.s. | 4.2E-07 |
| $D_v [mg/\sqrt{gl_{c0}}]$ | 3.4E-03 | n.s. | n.s. | n.s. | n.s. | 4.2E-18 | n.s. | n.s. | n.s. | n.s. |
| $x_{vpp} [l_0]$ | 2.4E-06 | n.s. | n.s. | n.s. | n.s. | 8.0E-15 | n.s. | n.s. | 1.8E-02 | n.s. |
| $z_{vpp} [l_0]$ | n.s. | n.s. | n.s. | 5.6E-03 | n.s. | 9.6E-24 | n.s. | 2.1E-03 | n.s. | n.s. |
| $xw_{vpp} [l_0]$ | 2.8E-02 | n.s. | n.s. | n.s. | n.s. | 1.9E-11 | n.s. | n.s. | n.s. | n.s. |
| $L_{cy} [m l_{c0} \sqrt{g l_{c0}}]$ | n.s. | n.s. | n.s. | n.s. | n.s. | 4.1E-07 | n.s. | n.s. | 1.9E-05 | n.s. |
| $W_{ax} [mg l_0]$ | n.s. | n.s. | n.s. | 4.2E-03 | 1.0E-02 | 1.8E-19 | n.s. | 1.3E-04 | n.s. | n.s. |
| $W_{tan} [mg l_0]$ | 2.6E-06 | 1.0E-04 | 1.3E-04 | n.s. | n.s. | 1.2E-14 | 7.1E-04 | n.s. | n.s. | 8.0E-06 |
| $W_{vax} [mg l_{c0}]$ | n.s. | n.s. | n.s. | 4.2E-04 | n.s. | 2.0E-21 | 7.6E-03 | 1.6E-03 | n.s. | 8.5E-04 |
| $W_{vtan} [mg l_{c0}]$ | 7.5E-08 | 2.8E-04 | n.s. | n.s. | n.s. | 3.6E-21 | n.s. | 1.8E-03 | n.s. | 2.9E-06 |
| $\Delta E_{kin,x} [mg l_{c0}]$ | n.s. | n.s. | n.s. | n.s. | n.s. | 2.0E-13 | n.s. | 1.1E-03 | n.s. | n.s. |
| $\Delta E_{kin,z} [mg l_{c0}]$ | 2.2E-02 | n.s. | n.s. | n.s. | n.s. | 1.3E-18 | n.s. | n.s. | n.s. | n.s. |
| $\Delta E_{pot} [mg l_{c0}]$ | 1.0E-07 | 1.7E-03 | 6.1E-03 | n.s. | n.s. | 1.3E-32 | 7.1E-04 | 1.0E-07 | 6.0E-07 | 3.7E-03 |
| $\Delta E_{ext} [mg l_{c0}]$ | 3.7E-08 | 3.1E-03 | 1.1E-03 | n.s. | n.s. | 1.9E-27 | n.s. | 1.3E-06 | 2.6E-02 | n.s. |

**Table S2.** Comparison of global parameters between the skipping and hurdling for the trailing and leading leg: timing, kinetics, leg properties, energetics. (Probabilities of univariate GLM)

|  | TRAILING |  |  | LEADING |  |  |
| --- | --- | --- | --- | --- | --- | --- |
|  | sk-hu | sk-hu*Fr | sk-hu*ani | sk-hu | sk-hu*Fr | sk-hu*ani |
| $T \left[ \sqrt{g/l_0} \right]$ | 7.3E-08 | 5.3E-09 | 1.0E-04 | 7.3E-08 | 5.3E-09 | 1.0E-04 |
| $t_c \left[ \sqrt{g/l_0} \right]$ | 1.9E-02 | 6.8E-15 | n.s. | n.s | 3.5E-19 | 1.4E-04 |
| $t_a \left[ \sqrt{g/l_0} \right]$ | 1.5E-14 | n.s. | 4.1E-05 | 1.3E-25 | 2.2E-13 | 1.3E-14 |
| $\beta_{leg-TD} (deg)$ | n.s. | n.s. | n.s. | 9.6E-16 | n.s. | n.s. |
| $\beta_{leg-LO} (deg)$ | 9.2E-05 | 1.5E-07 | 7.2E-05 | 1.2E-24 | 1.7E-02 | 7.4E-06 |
| $\beta_{lc-TD} (deg)$ | n.s. | n.s. | n.s. | 7.2E-08 | n.s. | n.s. |
| $\beta_{lc-LO} (deg)$ | n.s. | 3.6E-11 | 6.1E-06 | 1.4E-16 | n.s. | 4.4E-06 |
| $(l - l_0)_{TD} [l_0]$ | n.s. | n.s. | n.s. | n.s. | n.s. | n.s. |
| $(l - l_0)_{min} [l_0]$ | 6.6E-18 | n.s. | 7.1E-04 | n.s. | 1.9E-06 | 1.8E-06 |
| $(l - l_0)_{LO} [l_0]$ | n.s. | 8.8E-06 | n.s. | 2.7E-15 | 1.7E-13 | 1.0E-14 |
| $(l_c - l_{c0})_{TD} [l_{c0}]$ | n.s. | n.s. | n.s. | 4.0E-11 | 2.4E-08 | 9.3E-08 |
| $(l_c - l_{c0})_{min} [l_{c0}]$ | 2.5E-12 | n.s. | 5.4E-03 | 3.4E-13 | 5.2E-10 | 1.3E-05 |
| $(l_c - l_{c0})_{LO} [l_{c0}]$ | n.s. | 3.0E-03 | n.s. | 9.8E-17 | 4.5E-08 | 1.8E-07 |
| $x_{CoP1} [l_0]$ | n.s. | n.s. | n.s. | n.s. | n.s. | n.s. |
| $x_{CoP2} [l_0]$ | n.s. | 2.7E-02 | 3.0E-02 | n.s. | 1.1E-03 | 6.0E-08 |
| $\beta_{tru-TD}(deg)$ | 5.3E-05 | 8.8E-10 | 6.3E-13 | 1.3E-08 | 2.9E-10 | 1.0E-09 |
| $\beta_{tru-min}(deg)$ | 5.9E-07 | 1.1E-09 | 3.7E-09 | 1.0E-07 | 9.2E-09 | 2.7E-06 |
| $\beta_{tru-max}(deg)$ | 2.0E-07 | 5.4E-10 | 5.3E-12 | 1.4E-09 | 2.8E-13 | 4.5E-09 |
| $\beta_{tru-LO}(deg)$ | 4.8E-08 | 2.1E-10 | 1.9E-07 | 4.3E-10 | 1.7E-13 | 2.2E-08 |
| $F_{xa} [mg]$ | 5.3E-05 | 8.8E-10 | 6.3E-13 | 1.3E-08 | 2.9E-10 | 1.0E-09 |
| $F_{ym} [mg]$ | 5.9E-07 | 1.1E-09 | 3.7E-09 | 1.0E-07 | 9.2E-09 | 2.7E-06 |
| $F_z [mg]$ | 2.0E-07 | 5.4E-10 | 5.3E-12 | 1.4E-09 | 2.8E-13 | 4.5E-09 |
| *Skew* [ ] | 4.8E-08 | 2.1E-10 | 1.9E-07 | 4.3E-10 | 1.7E-13 | 2.2E-08 |
| *Kurtosis* [ ] | 7.1E-09 | n.s. | n.s. | n.s. | n.s. | n.s. |
| $p_x[m\sqrt{gl_{c0}}]$ | n.s. | n.s. | n.s. | n.s. | n.s. | n.s. |
| $p_{xa}[m\sqrt{gl_{c0}}]$ | n.s. | n.s. | n.s. | n.s. | n.s. | n.s. |
| $p_{xp}[m\sqrt{gl_{c0}}]$ | n.s. | n.s. | n.s. | n.s. | n.s. | n.s. |
| $p_y[m\sqrt{gl_{c0}}]$ | n.s. | 2.1E-02 | 3.3E-05 | n.s. | 2.0E-03 | 3.0E-08 |
| $p_{ym}[m\sqrt{gl_{c0}}]$ | n.s. | 5.4E-03 | 2.9E-06 | n.s. | 2.1E-03 | 3.4E-09 |
| $p_{yl}[m\sqrt{gl_{c0}}]$ | n.s. | n.s. | 2.1E-02 | n.s. | 2.0E-02 | 9.9E-04 |
| $p_z[m\sqrt{gl_{c0}}]$ | 7.1E-04 | n.s. | n.s. | 2.3E-09 | 5.6E-04 | 5.6E-04 |
| $p_z - mg t_c [m\sqrt{gl_{c0}}]$ | 1.7E-02 | 2.2E-06 | n.s. | 2.3E-11 | 9.5E-06 | 7.1E-04 |
| $k [mg/l_0]$ | 1.3E-07 | 3.0E-02 | 7.4E-06 | 6.8E-03 | n.s. | n.s. |
| $D [mg/\sqrt{gl_{c0}}]$ | n.s. | 4.5E-03 | n.s. | 4.0E-06 | n.s. | 2.5E-03 |
| $k_v [mg/l_{c0}]$ | 1.9E-06 | n.s. | 5.2E-06 | 3.4E-14 | n.s. | n.s. |
| $D_v [mg/\sqrt{gl_{c0}}]$ | 3.1E-08 | n.s. | n.s. | 5.6E-04 | n.s. | n.s. |
| $x_{vpp} [l_0]$ | n.s. | n.s. | n.s. | 3.2E-06 | n.s. | 2.8E-04 |
| $z_{vpp} [l_0]$ | 4.2E-04 | n.s. | n.s. | 1.5E-12 | 7.0E-05 | n.s. |
| $xw_{vpp} [l_0]$ | 4.6E-05 | n.s. | n.s. | 7.3E-05 | 2.3E-03 | 7.8E-06 |
| $L_{cy} [m l_{c0} \sqrt{g l_{c0}}]$ | n.s. | 1.3E-03 | n.s. | n.s. | 4.2E-04 | 9.0E-03 |
| $W_{ax} [mg l_0]$ | 9.5E-07 | 2.2E-02 | 5.5E-03 | 7.5E-15 | 1.8E-03 | 2.0E-06 |
| $W_{tan} [mg l_0]$ | n.s. | 2.8E-04 | 2.3E-05 | 6.4E-19 | 2.4E-03 | 4.0E-07 |
| $W_{vax} [mg l_{c0}]$ | 3.1E-05 | 9.3E-03 | n.s. | 7.4E-24 | 1.4E-04 | 1.3E-08 |
| $W_{vtan} [mg l_{c0}]$ | n.s. | 2.0E-08 | 4.7E-05 | 2.3E-11 | n.s. | 3.3E-07 |
| $\Delta E_{kin,x} [mg l_{c0}]$ | n.s. | n.s. | n.s. | n.s. | 1.1E+02 | 1.5E+00 |
| $\Delta E_{kin,z} [mg l_{c0}]$ | n.s. | n.s. | n.s. | 1.7E-10 | 1.3E-03 | 7.5E-05 |
| $\Delta E_{pot} [mg l_{c0}]$ | 3.3E-21 | 3.0E-11 | 2.9E-05 | 2.7E-28 | 2.3E-06 | 1.8E-11 |
| $\Delta E_{ext} [mg l_{c0}]$ | 2.2E-10 | 8.9E-03 | n.s. | 2.5E-21 | 3.5E-03 | 5.9E-10 |
| *Congruity* [ ] | 7.7E-15 | 4.4E-06 | n.s. | 7.7E-15 | 4.4E-06 | n.s. |
| *Recovery* [%] | 2.4E-03 | n.s. | n.s. | 2.4E-03 | n.s. | n.s. |
| *CoT* [mg] | 1.4E-20 | n.s. | 1.3E-07 | 1.4E-20 | n.s. | 1.3E-07 |

**Table S3.** Comparison of global parameters between the leading and trailing leg and skipping and hurdling: timing, kinetics, leg properties, energetics. (Probabilities of t- or Wilcoxon test)

| Variable | | SKIPPING | | | | | HURDLING | | | | | $p_{trlesk}$ | $p_{trlehu}^*$ |
| --- | --- | --- | --- | --- | --- | --- | --- | --- | --- | --- | --- | --- | --- |
| | | Mean | Std | Med | Min | Max | Mean | Std | Med | Min | Max | $p_{skhutr}$ | $p_{skhule}$ |
| $T \left[ \sqrt{g/l_0} \right]$ | stride | 2.416 | ±0.212 | 2.414 | 2.026 | 2.799 | 2.652 | ±0.185 | 2.653 | 2.197 | 3.094 | | 1.66E-03 |
| $t_c \left[ \sqrt{g/l_0} \right]$ | trail | 1.144 | ±0.146 | 1.095 | 0.970 | 1.441 | 1.218 | ±0.138 | 1.218 | 0.887 | 1.541 | n.s. | 3.80E-05 |
|  | lead | 1.102 | ±0.164 | 1.069 | 0.887 | 1.466 | 1.068 | ±0.067 | 1.069 | 0.970 | 1.268 | n.s. | n.s. |
| $t_a \left[ \sqrt{g/l_0} \right]$ | trail | -0.042 | ±0.043 | -0.039 | -0.139 | 0.000 | -0.350 | ±0.142 | -0.398 | -0.527 | 0.000 | 8.15E-05 | 1.30E-18 |
|  | lead | 0.298 | ±0.251 | 0.174 | 0.025 | 0.721 | 0.914 | ±0.235 | 0.870 | 0.531 | 1.580 | 3.43E-11 | 9.08E-11 |
| $\beta_{leg-TD} (deg)$ | trail | -27.07 | ±3.96 | -28.45 | -31.73 | -18.86 | -25.90 | ±4.13 | -25.45 | -36.73 | -19.46 | n.s. | 3.52E-06 |
|  | lead | -28.82 | ±2.11 | -29.08 | -31.87 | -25.56 | -37.26 | ±2.28 | -36.98 | -42.16 | -32.09 | n.s. | 2.30E-08 |
| $\beta_{leg-LO} (deg)$ | trail | 35.51 | ±4.80 | 35.95 | 25.65 | 43.69 | 31.50 | ±4.03 | 32.55 | 20.19 | 36.74 | 1.89E-03 | 3.52E-06 |
|  | lead | 27.43 | ±3.46 | 26.99 | 21.14 | 34.40 | 11.69 | ±3.42 | 10.95 | 5.62 | 20.01 | 1.14E-02 | 2.30E-08 |
| $\beta_{lc-TD} (deg)$ | trail | -17.08 | ±3.23 | -17.62 | -22.22 | -12.17 | -14.44 | ±3.22 | -13.48 | -20.66 | -9.85 | 1.89E-03 | 3.52E-06 |
|  | lead | -21.05 | ±1.86 | -21.12 | -23.64 | -18.12 | -25.84 | ±2.60 | -25.34 | -32.34 | -22.11 | 3.32E-02 | 4.38E-07 |
| $\beta_{lc-LO} (deg)$ | trail | 31.40 | ±3.85 | 31.88 | 24.08 | 37.71 | 30.83 | ±4.75 | 32.50 | 17.53 | 36.82 | 1.61E-03 | 1.08E-20 |
|  | lead | 23.59 | ±4.11 | 24.11 | 17.02 | 31.21 | 12.77 | ±3.61 | 13.24 | 5.92 | 18.68 | n.s. | 7.71E-08 |
| $(l-l_0)_{TD} [l_0]$ | trail | 0.972 | ±0.031 | 0.968 | 0.912 | 1.047 | 0.949 | ±0.025 | 0.944 | 0.911 | 1.005 | n.s. | 8.23E-05 |
|  | lead | 0.984 | ±0.028 | 0.981 | 0.939 | 1.039 | 0.979 | ±0.034 | 0.972 | 0.929 | 1.055 | 1.80E-02 | n.s. |
| $(l-l_0)_{min} [l_0]$ | trail | 0.859 | ±0.019 | 0.859 | 0.829 | 0.892 | 0.763 | ±0.029 | 0.760 | 0.715 | 0.833 | 1.13E-02 | 3.61E-17 |
|  | lead | 0.890 | ±0.030 | 0.889 | 0.838 | 0.930 | 0.883 | ±0.024 | 0.882 | 0.829 | 0.942 | 2.94E-08 | n.s. |
| $(l-l_0)_{LO} [l_0]$ | trail | 1.088 | ±0.083 | 1.071 | 0.980 | 1.286 | 1.063 | ±0.057 | 1.066 | 0.924 | 1.169 | n.s. | 3.52E-06 |
|  | lead | 1.073 | ±0.077 | 1.040 | 0.969 | 1.172 | 1.148 | ±0.040 | 1.138 | 1.087 | 1.264 | n.s. | 1.31E-04 |
| $(l_c-l_{c0})_{TD} [l_{c0}]$ | trail | 0.964 | ±0.027 | 0.961 | 0.917 | 1.007 | 0.952 | ±0.029 | 0.951 | 0.901 | 1.045 | n.s. | 5.46E-04 |
|  | lead | 0.977 | ±0.028 | 0.974 | 0.918 | 1.033 | 0.917 | ±0.047 | 0.899 | 0.852 | 1.011 | n.s. | 3.03E-04 |
| $(l_c-l_{c0})_{min} [l_{c0}]$ | trail | 0.900 | ±0.015 | 0.903 | 0.873 | 0.933 | 0.836 | ±0.029 | 0.834 | 0.788 | 0.897 | 2.21E-03 | 2.37E-05 |
|  | lead | 0.926 | ±0.019 | 0.931 | 0.887 | 0.949 | 0.869 | ±0.039 | 0.860 | 0.799 | 0.939 | 1.39E-07 | 3.46E-05 |
| $(l_c-l_{c0})_{LO} [l_{c0}]$ | trail | 1.073 | ±0.048 | 1.058 | 1.010 | 1.193 | 1.068 | ±0.042 | 1.082 | 0.976 | 1.133 | n.s. | 3.52E-06 |
|  | lead | 1.070 | ±0.038 | 1.072 | 1.021 | 1.123 | 1.134 | ±0.021 | 1.132 | 1.108 | 1.201 | n.s. | 3.12E-07 |
| $x_{CoP1} [l_0]$ | trail | 0.043 | ±0.027 | 0.041 | 0.005 | 0.097 | 0.039 | ±0.029 | 0.034 | 0.000 | 0.095 | 4.16E-02 | n.s. |
|  | lead | 0.024 | ±0.020 | 0.017 | 0.003 | 0.073 | 0.039 | ±0.032 | 0.035 | 0.001 | 0.154 | n.s. | n.s. |
| $x_{CoP2} [l_0]$ | trail | 0.148 | ±0.052 | 0.147 | 0.049 | 0.236 | 0.118 | ±0.056 | 0.103 | 0.040 | 0.275 | n.s. | 3.52E-06 |
|  | lead | 0.179 | ±0.032 | 0.180 | 0.124 | 0.222 | 0.201 | ±0.046 | 0.183 | 0.147 | 0.317 | n.s. | n.s. |
| $\beta_{tru-TD}(deg)$ | trail | 29.04 | ±5.90 | 27.10 | 39.76 | 19.23 | 32.91 | ±5.75 | 35.04 | 40.20 | 20.83 | n.s. | 3.88E-05 |
|  | lead | 28.20 | ±4.82 | 28.72 | 35.62 | 19.64 | 35.59 | ±7.99 | 39.03 | 46.08 | 19.74 | n.s. | 4.06E-03 |
| $\beta_{tru-min}(deg)$ | trail | 30.07 | ±5.99 | 28.86 | 40.21 | 20.67 | 36.72 | ±7.58 | 39.55 | 50.25 | 21.84 | n.s. | n.s. |
|  | lead | 28.85 | ±4.69 | 30.26 | 36.28 | 21.45 | 36.63 | ±7.71 | 39.54 | 50.52 | 21.11 | 5.03E-03 | 2.43E-03 |
| $\beta_{tru-max}(deg)$ | trail | 26.79 | ±5.09 | 25.50 | 35.43 | 19.10 | 32.18 | ±6.62 | 34.85 | 39.29 | 19.33 | 1.71E-02 | n.s. |
|  | lead | 25.22 | ±4.76 | 27.09 | 32.98 | 17.73 | 32.53 | ±7.91 | 36.18 | 43.13 | 17.80 | 7.14E-03 | 3.27E-03 |
| $\beta_{tru-LO}(deg)$ | trail | 27.69 | ±5.19 | 27.02 | 36.15 | 20.05 | 35.75 | ±8.51 | 38.49 | 50.25 | 19.33 | 1.89E-03 | 6.70E-09 |
|  | lead | 25.30 | ±4.76 | 27.22 | 33.31 | 18.32 | 32.75 | ±7.60 | 36.18 | 43.13 | 18.55 | 2.82E-03 | 2.43E-03 |
| $F_{xa} [mg]$ | trail | 0.213 | ±0.040 | 0.209 | 0.148 | 0.298 | 0.250 | ±0.029 | 0.253 | 0.178 | 0.304 | n.s. | 3.64E-02 |
|  | lead | 0.218 | ±0.053 | 0.213 | 0.141 | 0.380 | 0.271 | ±0.031 | 0.275 | 0.206 | 0.329 | 6.22E-03 | 8.02E-05 |
| $F_{ym} [mg]$ | trail | -0.170 | ±0.055 | -0.164 | -0.260 | -0.072 | -0.155 | ±0.047 | -0.151 | -0.298 | -0.089 | n.s. | n.s. |
|  | lead | -0.138 | ±0.093 | -0.113 | -0.300 | -0.006 | -0.163 | ±0.058 | -0.142 | -0.320 | -0.069 | n.s. | n.s. |
| $F_z$ [mg] | trail | 1.618 | ±0.148 | 1.581 | 1.401 | 1.905 | 1.927 | ±0.270 | 1.817 | 1.594 | 2.606 | n.s. | n.s. |
|  | lead | 1.571 | ±0.224 | 1.648 | 1.155 | 1.875 | 1.892 | ±0.163 | 1.939 | 1.610 | 2.211 | 6.67E-05 | 6.67E-05 |
| Skew [ ] | trail | 0.312 | ±0.063 | 0.306 | 0.190 | 0.417 | 0.398 | ±0.058 | 0.405 | 0.233 | 0.510 | 3.68E-02 | 3.52E-06 |
|  | lead | 0.268 | ±0.066 | 0.243 | 0.158 | 0.391 | 0.179 | ±0.056 | 0.187 | 0.055 | 0.258 | 1.15E-04 | 3.03E-04 |
| Kurtosis [ ] | trail | -0.703 | ±0.068 | -0.695 | -0.860 | -0.593 | -0.490 | ±0.113 | -0.457 | -0.717 | -0.325 | 1.71E-02 | 2.74E-15 |
|  | lead | -0.784 | ±0.083 | -0.789 | -0.911 | -0.628 | -0.834 | ±0.049 | -0.834 | -0.942 | -0.736 | 1.33E-06 | n.s. |
| $p_x[m\sqrt{gl_{c0}}]$ | trail | -0.005 | ±0.032 | -0.001 | -0.084 | 0.057 | 0.013 | ±0.030 | 0.010 | -0.045 | 0.085 | n.s. | n.s. |
|  | lead | 0.006 | ±0.040 | -0.006 | -0.048 | 0.119 | 0.017 | ±0.028 | 0.020 | -0.056 | 0.083 | n.s. | n.s. |
| $p_{xa}[m\sqrt{gl_{c0}}]$ | trail | 0.083 | ±0.017 | 0.079 | 0.055 | 0.114 | 0.102 | ±0.024 | 0.096 | 0.061 | 0.162 | n.s. | n.s. |
|  | lead | 0.086 | ±0.025 | 0.083 | 0.053 | 0.156 | 0.102 | ±0.017 | 0.098 | 0.083 | 0.165 | 2.31E-02 | 8.76E-03 |
| $p_{xp}[m\sqrt{gl_{c0}}]$ | trail | -0.088 | ±0.020 | -0.084 | -0.140 | -0.057 | -0.089 | ±0.018 | -0.089 | -0.141 | -0.051 | n.s. | n.s. |
|  | lead | -0.080 | ±0.018 | -0.087 | -0.102 | -0.037 | -0.085 | ±0.019 | -0.084 | -0.138 | -0.048 | n.s. | n.s. |
| $p_y[m\sqrt{gl_{c0}}]$ | trail | 0.045 | ±0.044 | 0.028 | -0.014 | 0.107 | 0.040 | ±0.021 | 0.042 | -0.008 | 0.075 | n.s. | n.s. |
|  | lead | 0.034 | ±0.061 | 0.032 | -0.049 | 0.158 | 0.044 | ±0.039 | 0.034 | -0.005 | 0.132 | n.s. | n.s. |
| $p_{ym}[m\sqrt{gl_{c0}}]$ | trail | 0.063 | ±0.035 | 0.050 | 0.021 | 0.116 | 0.061 | ±0.016 | 0.065 | 0.027 | 0.096 | n.s. | n.s. |
|  | lead | 0.054 | ±0.047 | 0.046 | 0.000 | 0.161 | 0.060 | ±0.032 | 0.046 | 0.024 | 0.134 | n.s. | n.s. |
| $p_{yl}[m\sqrt{gl_{c0}}]$ | trail | -0.017 | ±0.009 | -0.017 | -0.035 | -0.008 | -0.022 | ±0.008 | -0.022 | -0.042 | -0.003 | n.s. | 7.16E-03 |
|  | lead | -0.020 | ±0.015 | -0.015 | -0.049 | -0.002 | -0.015 | ±0.009 | -0.015 | -0.035 | -0.001 | n.s. | n.s. |
| $p_z[m\sqrt{gl_{c0}}]$ | trail | 1.156 | ±0.129 | 1.119 | 0.897 | 1.412 | 1.342 | ±0.143 | 1.285 | 1.101 | 1.613 | n.s. | n.s. |
|  | lead | 1.120 | ±0.141 | 1.152 | 0.938 | 1.297 | 1.334 | ±0.115 | 1.338 | 1.110 | 1.638 | 3.90E-04 | 2.59E-05 |
| $p_z - mg t_c$<br>[$m\sqrt{gl_{c0}}$] | trail | 0.017 | ±0.161 | 0.025 | -0.214 | 0.296 | 0.234 | ±0.304 | 0.144 | -0.195 | 0.897 | n.s. | 1.62E-02 |
|  | lead | 0.023 | ±0.353 | 0.015 | -0.487 | 0.509 | 0.535 | ±0.213 | 0.571 | 0.158 | 0.884 | 3.32E-02 | 3.46E-05 |
| $k$ [mg/$l_0$] | trail | 9.688 | ±2.166 | 9.209 | 6.347 | 14.071 | 7.156 | ±1.511 | 6.657 | 5.398 | 11.526 | 1.71E-02 | 1.97E-05 |
|  | lead | 11.822 | ±1.260 | 11.708 | 9.009 | 14.673 | 10.064 | ±1.528 | 10.311 | 6.596 | 12.019 | 1.05E-04 | 1.22E-03 |
| $D$ [mg/$\sqrt{gl_{c0}}$] | trail | 0.000 | ±0.441 | -0.008 | -1.034 | 0.732 | 0.246 | ±0.212 | 0.253 | -0.350 | 0.732 | 9.81E-04 | 3.52E-06 |
|  | lead | -0.524 | ±0.658 | -0.324 | -1.666 | 0.357 | -1.290 | ±0.447 | -1.389 | -2.040 | -0.243 | n.s. | 8.19E-04 |
| $k_v$ [mg/$l_{c0}$] | trail | 14.123 | ±3.149 | 13.788 | 8.589 | 18.727 | 10.338 | ±2.842 | 9.857 | 7.399 | 17.942 | n.s. | n.s. |
|  | lead | 16.528 | ±2.293 | 16.037 | 11.484 | 21.100 | 9.811 | ±1.565 | 9.995 | 7.335 | 13.073 | 3.03E-04 | 3.75E-08 |
| $D_v$ [mg/$\sqrt{gl_{c0}}$] | trail | -0.682 | ±0.397 | -0.764 | -1.217 | 0.039 | -0.034 | ±0.277 | -0.047 | -0.814 | 0.554 | 2.21E-03 | 3.52E-06 |
|  | lead | -1.463 | ±0.685 | -1.579 | -2.803 | -0.266 | -2.313 | ±0.568 | -2.506 | -3.117 | -0.966 | 1.07E-05 | 3.03E-04 |
| $x_{vpp}$ [$l_0$] | trail | -0.108 | ±0.048 | -0.123 | -0.193 | -0.004 | -0.162 | ±0.069 | -0.173 | -0.286 | -0.042 | 9.81E-04 | 3.52E-06 |
|  | lead | 0.035 | ±0.052 | 0.042 | -0.069 | 0.132 | 0.111 | ±0.057 | 0.096 | 0.025 | 0.215 | n.s. | 5.91E-04 |
| $z_{vpp}$ [$l_0$] | trail | 0.270 | ±0.146 | 0.257 | 0.032 | 0.542 | 0.432 | ±0.094 | 0.432 | 0.260 | 0.642 | 9.86E-03 | 3.52E-06 |
|  | lead | 0.090 | ±0.177 | 0.040 | -0.104 | 0.612 | -0.205 | ±0.076 | -0.211 | -0.336 | -0.011 | 7.55E-04 | 9.78E-08 |
| $xw_{vpp}$ [$l_0$] | trail | 0.190 | ±0.043 | 0.196 | 0.111 | 0.266 | 0.329 | ±0.105 | 0.336 | 0.141 | 0.516 | 2.59E-03 | 3.52E-06 |
|  | lead | 0.138 | ±0.054 | 0.146 | 0.063 | 0.219 | 0.099 | ±0.024 | 0.094 | 0.064 | 0.152 | 5.54E-05 | n.s. |
| $L_{cy}$ [$m l_{c0} \sqrt{gl_{c0}}$] | trail | -0.171 | ±0.287 | -0.022 | -0.822 | 0.108 | -0.114 | ±0.139 | -0.157 | -0.338 | 0.181 | n.s. | 2.60E-05 |
|  | lead | 0.092 | ±0.271 | 0.037 | -0.201 | 1.047 | 0.171 | ±0.206 | 0.108 | -0.106 | 0.753 | n.s. | n.s. |
| $W_{ax}$ [mg $l_0$] | trail | 0.019 | ±0.071 | 0.001 | -0.085 | 0.157 | -0.081 | ±0.064 | -0.076 | -0.214 | 0.067 | 7.42E-03 | 3.52E-06 |
|  | lead | 0.063 | ±0.078 | 0.035 | -0.037 | 0.190 | 0.211 | ±0.053 | 0.206 | 0.095 | 0.306 | 5.54E-05 | 1.19E-06 |
| $W_{tan}$ [mg $l_0$] | trail | 0.005 | ±0.080 | 0.010 | -0.131 | 0.145 | 0.054 | ±0.070 | 0.028 | -0.043 | 0.260 | 2.87E-02 | 2.68E-12 |
|  | lead | 0.082 | ±0.029 | 0.075 | 0.048 | 0.163 | 0.246 | ±0.066 | 0.268 | 0.085 | 0.373 | n.s. | 8.62E-13 |
| $W_{vax}$ [mg $l_{c0}$] | trail | 0.065 | ±0.042 | 0.063 | -0.003 | 0.133 | 0.003 | ±0.047 | 0.012 | -0.152 | 0.105 | n.s. | 3.52E-06 |
|  | lead | 0.086 | ±0.041 | 0.081 | 0.017 | 0.156 | 0.293 | ±0.064 | 0.306 | 0.135 | 0.394 | 1.38E-04 | 3.32E-08 |
| $W_{vtan} [mg l_{c0}]$ | trail | -0.069 | $\pm 0.045$ | -0.079 | -0.132 | 0.006 | -0.080 | $\pm 0.033$ | -0.092 | -0.118 | -0.008 | 7.00E-04 | 3.52E-06 |
| | lead | 0.023 | $\pm 0.034$ | 0.023 | -0.044 | 0.086 | 0.088 | $\pm 0.034$ | 0.083 | 0.019 | 0.140 | n.s. | 1.44E-05 |
| $\Delta E_{kin,x} [mg l_{c0}]$ | trail | -0.024 | $\pm 0.025$ | -0.019 | -0.082 | 0.016 | -0.041 | $\pm 0.030$ | -0.045 | -0.095 | 0.016 | 4.11E-03 | 5.75E-06 |
| | lead | 0.007 | $\pm 0.034$ | 0.000 | -0.044 | 0.112 | 0.035 | $\pm 0.027$ | 0.035 | -0.038 | 0.083 | n.s. | 1.66E-03 |
| $\Delta E_{kin,z} [mg l_{c0}]$ | trail | -0.016 | $\pm 0.013$ | -0.012 | -0.038 | 0.001 | -0.041 | $\pm 0.036$ | -0.049 | -0.138 | 0.085 | 1.16E-03 | 5.33E-19 |
| | lead | 0.005 | $\pm 0.012$ | 0.001 | -0.008 | 0.036 | 0.062 | $\pm 0.038$ | 0.052 | 0.009 | 0.180 | 9.62E-05 | 4.42E-07 |
| $\Delta E_{pot} [mg l_{c0}]$ | trail | -0.051 | $\pm 0.036$ | -0.039 | -0.120 | 0.004 | -0.158 | $\pm 0.040$ | -0.169 | -0.228 | -0.071 | 8.29E-04 | 3.52E-06 |
| | lead | 0.062 | $\pm 0.045$ | 0.046 | -0.006 | 0.133 | 0.221 | $\pm 0.037$ | 0.218 | 0.128 | 0.294 | 1.56E-07 | 2.94E-08 |
| $\Delta E_{ext} [mg l_{c0}]$ | trail | -0.091 | $\pm 0.065$ | -0.085 | -0.206 | 0.001 | -0.240 | $\pm 0.068$ | -0.251 | -0.338 | -0.022 | 5.89E-04 | 3.52E-06 |
| | lead | 0.074 | $\pm 0.058$ | 0.064 | -0.001 | 0.189 | 0.318 | $\pm 0.091$ | 0.304 | 0.114 | 0.544 | 2.27E-06 | 8.68E-08 |
| *Congruity* [ ] | stride | 0.709 | $\pm 0.140$ | 0.755 | 0.449 | 0.878 | 0.481 | $\pm 0.038$ | 0.468 | 0.421 | 0.600 | | 9.90E-11 |
| *Recovery* [%] | stride | 5.646 | $\pm 4.197$ | 4.317 | 0.313 | 13.828 | 10.030 | $\pm 3.262$ | 10.344 | 2.432 | 13.911 | | 3.04E-03 |
| *CoT* [mg] | stride | 0.117 | $\pm 0.021$ | 0.117 | 0.086 | 0.165 | 0.195 | $\pm 0.024$ | 0.195 | 0.143 | 0.254 | | 3.75E-08 |

### FIGURES SUPPLEMENT

**Fig. S1.**
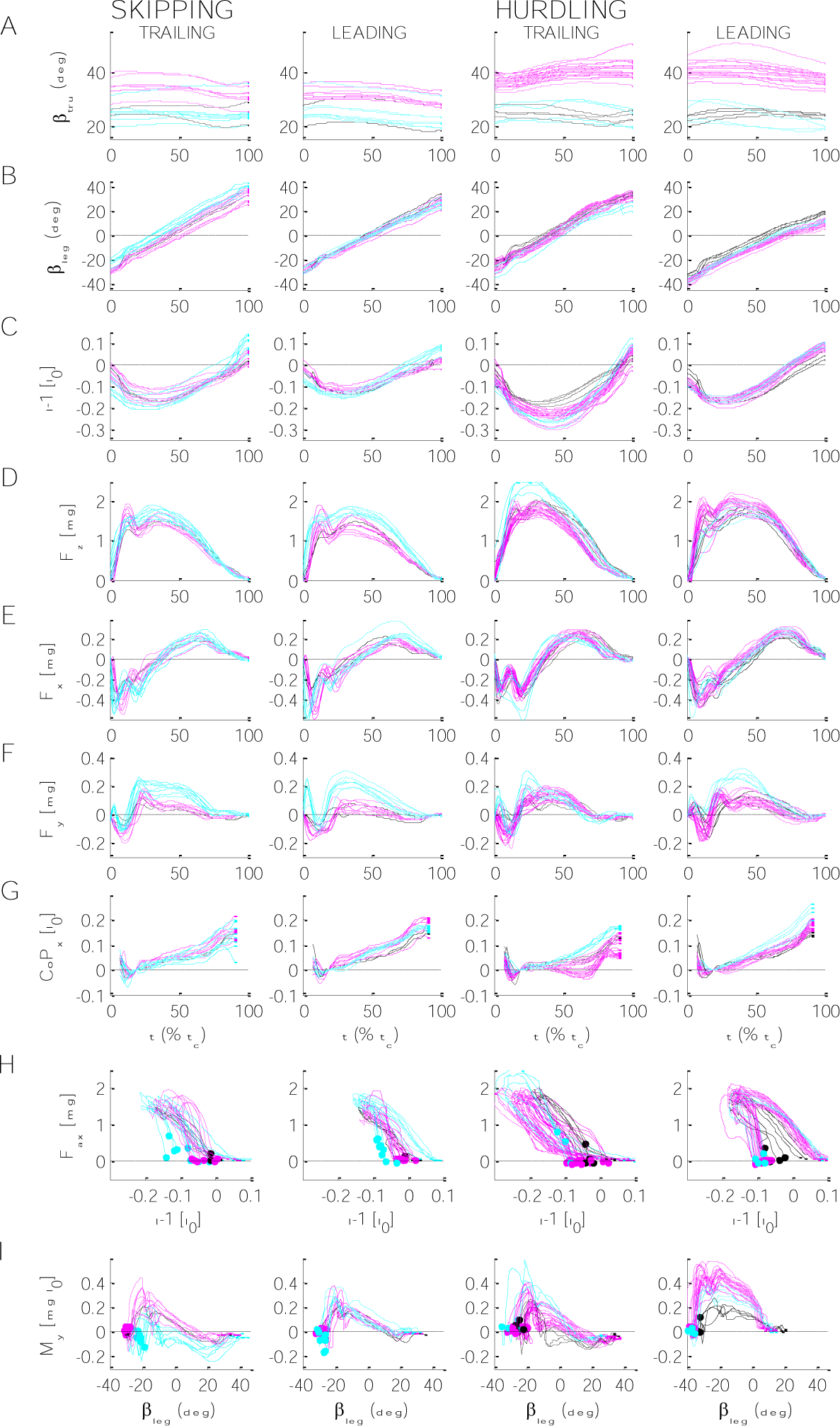
Individual global properties during skipping and hurdling in the trailing and the leading leg. (A-G) Time courses. *t*, time normalized to contact time, *t*_*c*_. (A) Trunk pitch, β_*tru*_. (B) Leg angle, β_*leg*_. (C) Change of leg length, (*l* − *l*_0_) *l*_0−1_. (D, E, F) Craniad, *F*_*z*_, anteriad, *F*_*x*_, and mediad, *F*_*y*_, components of ground reaction force. (G) Anteriad component of center of pressure, *CoP*_*x*_. *CoP*_*x*_ at 20% *t*_*c*_ set to 0 and the first and last 5% are omitted. H) Axial force length loops, *F*_*ax*_(*l* − *l*_0_)*l*_0−1_. Green dashed lines: fittings based on Voigt-model. Filled circles: touch down. I) Tangential moment angle loops, *M*_*y*_(β_*leg*_). Filled circles: touch down. Right leg: solid lines; left leg: dashed; macaques: black, Ku, magenta, Fu, cyan, Po.

**Fig. S2.**
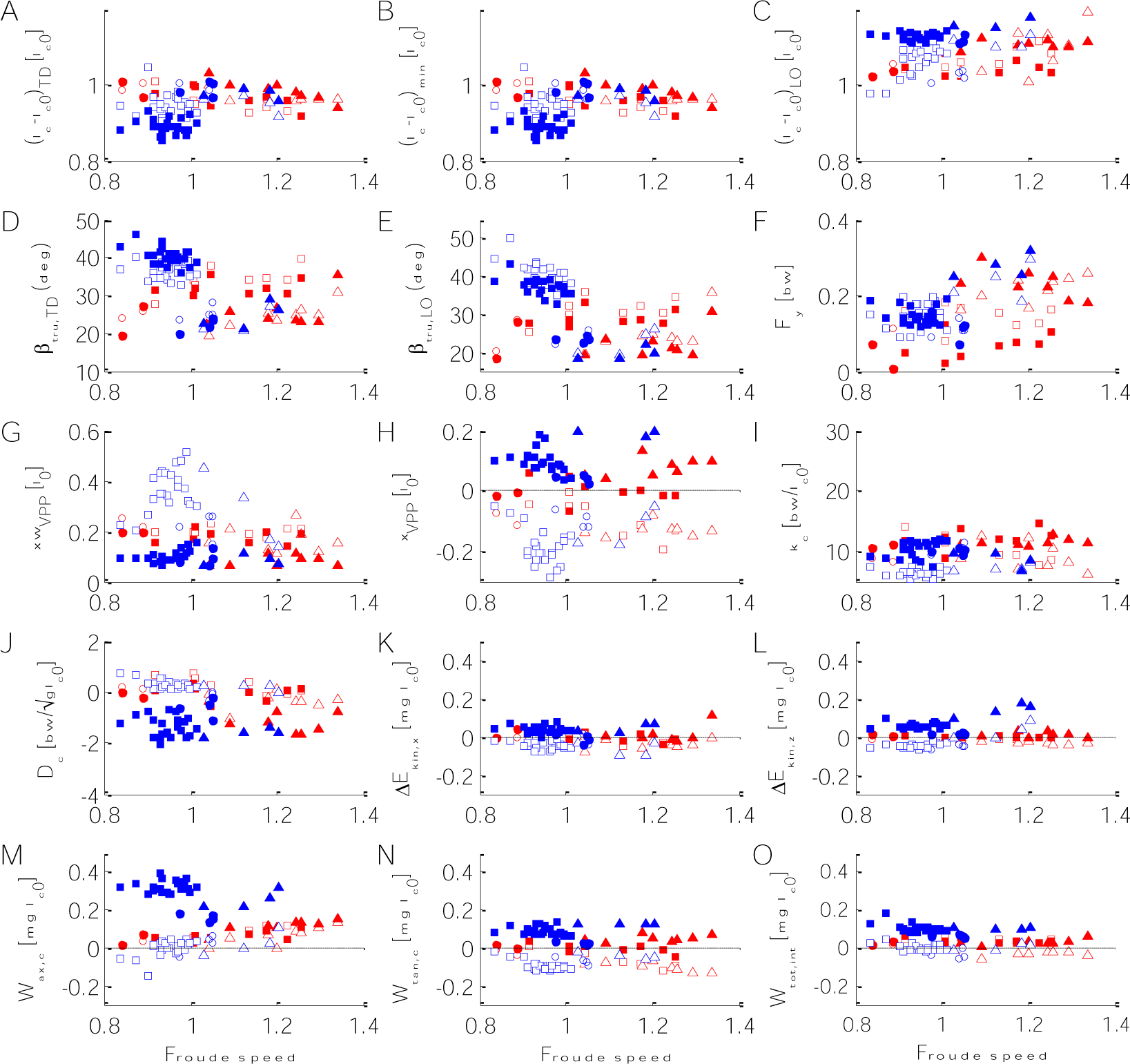
Dependence of kinematic, dynamic, and energetic stance parameters on Froude speed. Red: skipping; blue: hurdling; open: trailing leg; filled: leading leg. (A-C) Lengths of the virtual leg from CoM to CoP: (A) at touch down, (*l*_*c*_ − *l*_*c*0_)_*TD*_; (E) at minimum length, (*l*_*c*_ − *l*_*c*0_)_*min*_; (F) and lift off (*l*_*c*_ − *l*_*c*0_)_*LO*_. (D,E) Trunk angle with respect to the vertical: (D) at touch down, β_*tru,TD*_; (D) at lift off, β_*tru,LO*_. (F) Amplitude of the medial ground reaction force, *F*_*y*_. (G) Width of the VPP, *xw*_*VPP*_. (H) Anterior shift of the VPP, *x*_*VPP*_. (I) Stiffness of the virtual leg, *k*_*c*_. (J) Damping of the virtual leg, *D*_*c*_. (K,L) changes in kinetic energy of the CoM: (K) anterior, Δ*E*_*kin,x*_; (L) vertical, Δ*E*_*kin,z*_. (M,N) Work of the virtual leg: (M) axial work, *W*_*ax,c*_ ; (N) tangential work, *W*_*tan,c*_ ; (O) Internal work, *W*_*tot,int*_. Circle: Ku; square: Fu; triangle: Po.

**Fig. S3.**
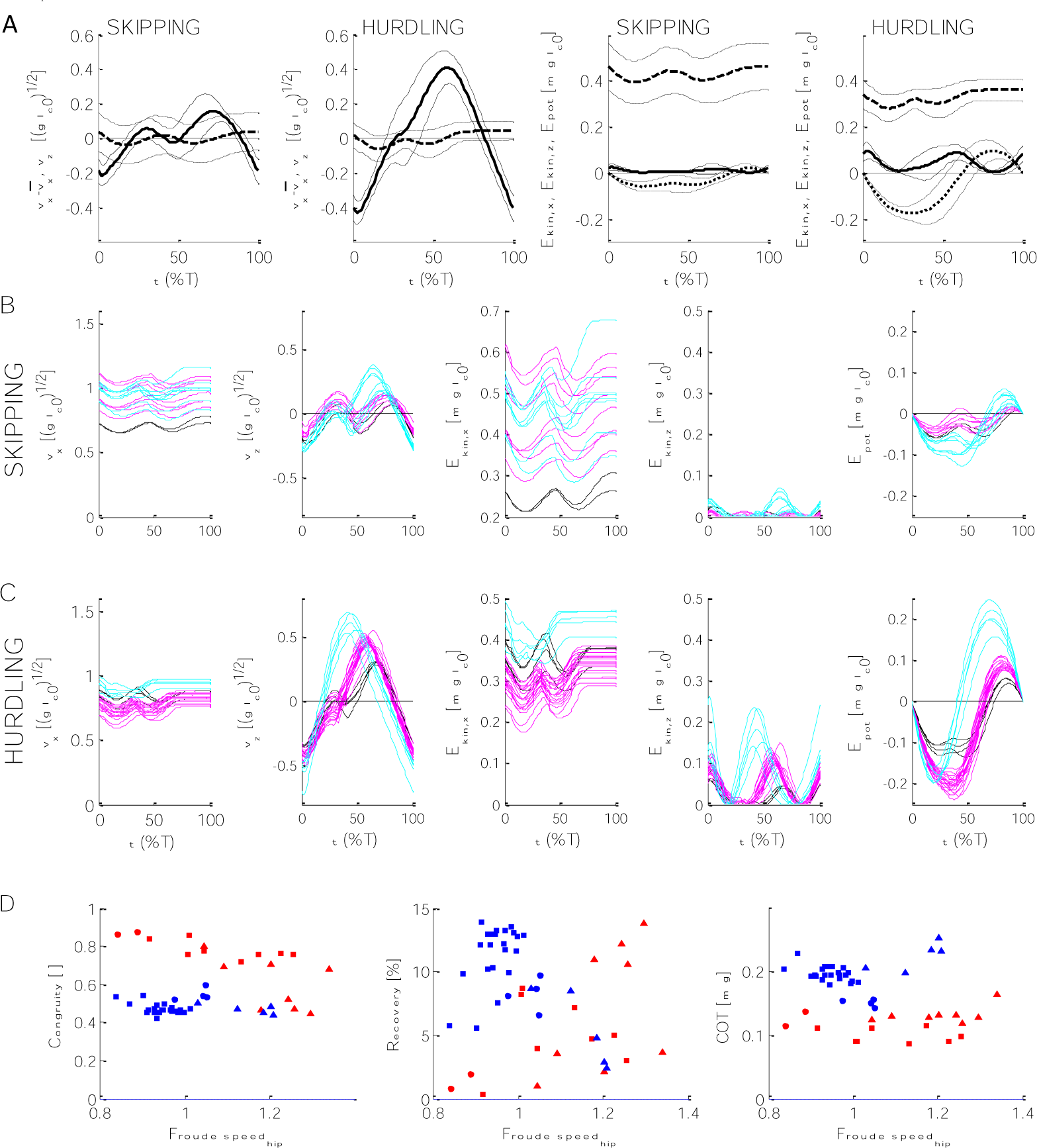
Velocity and energetics of the CoM during skipping and hurdling. A) Mean ±SD of the time courses of (from left to right) antorad velocity *v*_*x*_ minus its mean during the stride 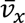 (dashed lines) as well as the vertical velocity *v*_*z*_(solid lines) for skipping and for hurdling. The kinetic energies *E*_*kin,x*_ (dashed lines), *E*_*kin,z*_ (solid lines), and potential energy *E*_*pot*_ (dotted lines) for skipping and hurdling. B,C) For skipping (B) and hurdling (C) for *v*_*x,z*_ and for *E*_*kin,x,z*_ and *E*_*pot*_ the tracings for each trial. Macaques: Ku - black; Fu – magenta; Po - cyan. D) From left to right: Congruity, recovery, and cost of transport (COT) during skipping (blue) and hurdling (red). Circle: Ku; square: Fu; triangle: Po.

